# A unified model of the task-evoked pupil response

**DOI:** 10.1101/2021.04.09.439231

**Authors:** Charlie S. Burlingham, Saghar Mirbagheri, David J. Heeger

**Affiliations:** Department of Psychology, New York University, New York, NY, 10003; Graduate Program in Neuroscience, University of Washington, Seattle, WA, 98195; Department of Psychology, Center for Neural Science, New York University, New York, NY, 10003

**Keywords:** arousal, cognition, task-evoked pupil response, oculomotor suppression, saccade, microsaccade

## Abstract

The pupil dilates and re-constricts following task events. It is popular to model this task-evoked pupil response as a linear transformation of event-locked impulses, the amplitudes of which are used as estimates of arousal. We show that this model is incorrect, and we propose an alternative model based on the physiological finding that a common neural input drives saccades and pupil size. The estimates of arousal from our model agreed with key predictions: arousal scaled with task difficulty and behavioral performance but was invariant to trial duration. Moreover, the model offers a unified explanation for a wide range of phenomena: entrainment of pupil size and saccade occurrence to task timing, modulation of pupil response amplitude and noise with task difficulty, reaction-time dependent modulation of pupil response timing and amplitude, a constrictory pupil response time-locked to saccades, and task-dependent distortion of this saccade-locked pupil response.

## 1 Introduction

Although the pupil most dramatically responds to changes in luminance [1, 2] and accommodation [1, 3, 4, 5], it fluctuates in size even in their absence [6]. During fixation and in the absence of a task, pupil size changes constantly and in a seemingly random way [6, 7] (Fig. S1A). During task performance, pupil size entrains to trial timing, [8, 9], dilating and then constricting sluggishly following trial onsets [8, 10, 11, 12] (Fig. S1C). This so-called “task-evoked” pupil response is modulated in amplitude by task demands, behavioral performance, and surprise [13, 14, 15, 16, 17, 18, 19, 20, 21]. Pupil responses covary with sweating and cardiac activity [22, 23], measures of peripheral autonomic activity, as well as spiking activity in the locus coeruleus [24, 25, 26, 27], basal forebrain [24, 28, 25], and dorsal raphé [20], sources of cortical norepinephrine, acetylcholine, and serotonin, respectively. Based on these observations, many studies have used pupil size as a non-invasive measure of arousal level [29, 30, 31, 32, 33].

The task-evoked pupil response is commonly used to estimate arousal by assuming pupil size is a linear transformation of task events (e.g., stimulus onset, button press, feedback). Specifically, existing models (Fig. 1A) assume that pupil size is the output of a dilatory low-pass filter, which operates on a series of impulses aligned with task events [34, 35, 36, 37, 38, 39, 40, 41, 42]. Arousal level, estimated as the amplitude of these inputs, is estimated using linear regression. We will refer to this approach as the “consensus model,” acknowledging that there are important predecessors in the literature [43, 44]. Although the consensus model is widely used, its central assumption — that pupil size is a linear transformation of task events — has not been tested.

**Figure 1:**
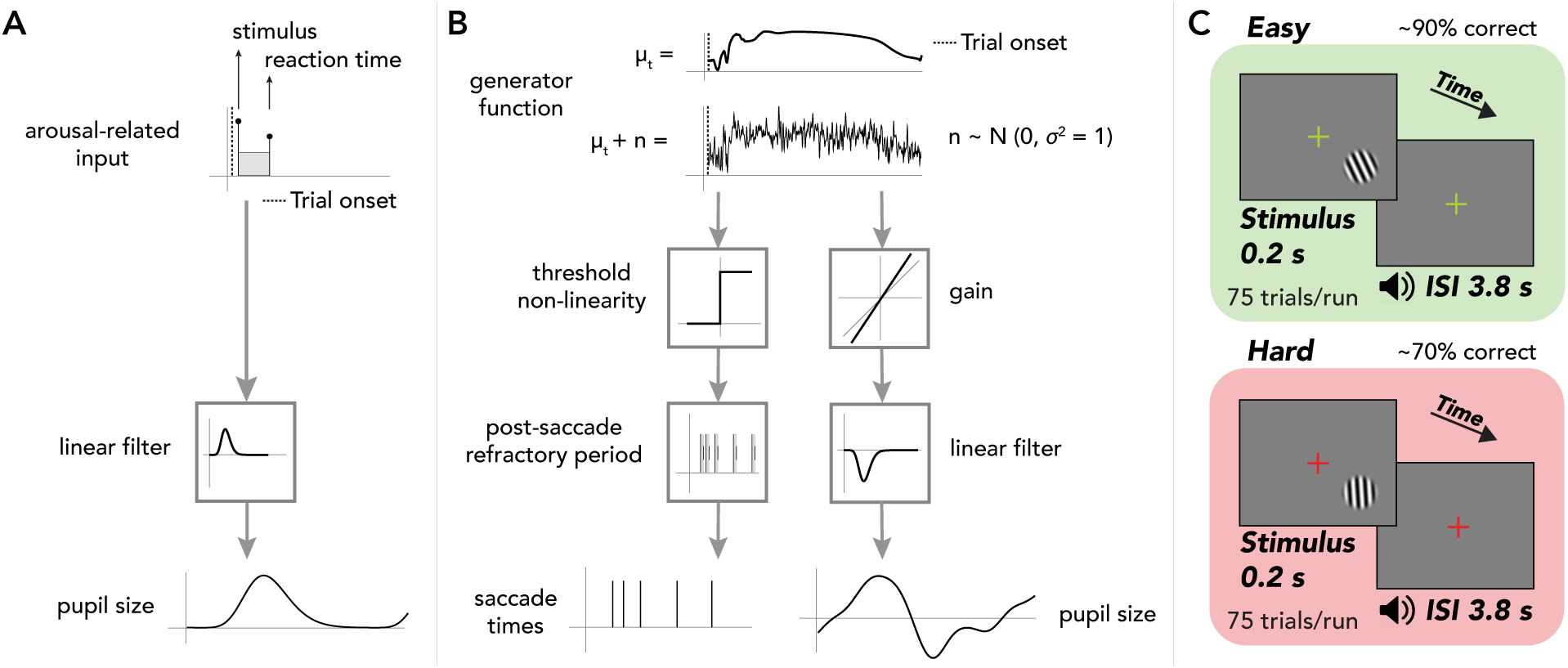
Models of the task-evoked pupil response and task protocol. (**A**) Consensus model. Arousal-related input (impulses at stimulus onset and reaction time plus a time-on-task boxcar) is filtered by a dilatory low-pass filter, predicting pupil size on a single trial [34, 35, 36, 38, 42]. (**B**) Linear-nonlinear linking model. The generator function, with normally-distributed noise (*n*) added to its expected value (*µ_t_*), is (1) subjected to a threshold and post-saccade refractory period, giving rise to a sequence of saccades, and (2) multiplied by a gain and filtered by a constrictory low-pass filter, producing pupil size. Generator function, *σ* reduced by 3x, Gaussian-blurred for visualization. Refractory period, grey regions. (**C**) Task, orientation discrimination around vertical. Timing, 0.2 s stimulus presentation, followed by a 3.8 s inter-stimulus interval (ISI). Observer had to respond during ISI with a button press and immediately received auditory feedback (correct, incorrect). Design, alternation between separate easy (approx. 90% correct) and hard (approx. 70% correct) runs, 75 trials each. Fixation cross, green or red, indicating easy or hard difficulty. Stimulus, grating (enlarged for visualization).

We tested the consensus model with a visual orientation-discrimination experiment in humans that varied in difficulty and trial duration. Estimates of arousal from the consensus model were volatile across tasks with different trial durations, demonstrating that pupil size is not a linear transformation of task events.

We propose an alternative model linking the task-evoked pupil response and arousal, which leverages the “common drive hypothesis” — i.e., that pupil size and saccades are driven by a common neural input — to constrain estimation of arousal [45, 46, 47, 10, 26, 21].

There is substantial support for the common drive hypothesis. A transient pupil constriction and re-dilation locked to saccades was reported in humans as early as the 1960s [48, 49, 1, 50, 51, 52, 53]. Microstimulation of superior colliculus (SC), a subcortical center for saccade generation, evokes a transient pupil response above or below the current level required to evoke a saccade [46, 26, 45]. Furthermore, converging anatomical [10, 2, 54], physiological [55, 1, 10, 26], and pharmacological evidence [56, 57, 58, 59, 6] supports predominantly parasympathetic control of the task-evoked pupil response via the preganglionic Edinger-Westphal nucleus (EW), which receives direct projections from the SC [10]. Involuntary fixational saccades (including microsaccades in our definition), like pupil responses, fluctuate with engagement — entraining to task timing. Indeed the two are correlated in a wide array of tasks [60, 17, 61, 62, 63, 64, 65, 66, 12, 13, 67]. Specifically, saccade occurrence is suppressed nearly completely at trial onset, resumes quickly thereafter, and then is more slowly suppressed again in anticipation of the next trial [68, 69, 70, 71, 67, 60, 72]. The magnitude of this oculomotor suppression has been linked to temporal attention [60, 73], temporal expectation [15, 72, 73], and task difficulty [67, 74, 75], task variables that have also been linked to pupil responses [13, 44, 17, 76, 42].

Our linking model has two channels (Fig. 1B). The first relates the common input to saccades and the second relates the input to pupil size. In the first channel, a saccade is generated whenever a noisy generator function crosses a fixed threshold. In the second channel, the same generator function is amplified by a gain and low-pass filtered, yielding pupil size. We offer an algorithm, based on this model, that estimates the gain and generator function from measurements of pupil size and saccade rate.

Our results strongly support the hypothesis that pupil size and saccades are driven by a common input [10, 45, 21] and extends that hypothesis to account for the influence of arousal on the task-evoked pupil response. Saccade timing, transformed with our model, accurately predicted the task-evoked pupil response. Gain, our model’s estimate of arousal, scaled with task difficulty and behavioral performance and generalized across different tasks. The generator function was modulated by the timing of the task and behavioral response, suggesting that the input to the pupil reflects a cognitive representation of task structure. Our model offers a unified explanation for a wide range of phenomena, including *(i)* entrainment of pupil size and saccade occurrence to task timing, *(ii)* modulation of pupil response amplitude with task difficulty *(iii)* modulation of trial-to-trial pupil noise with task difficulty, *(iv)* reaction-time dependent modulation of pupil response timing, *(v)* a constrictory pupil response time-locked to saccades, and *(vi)* task-dependent distortion of this saccade-locked pupil response.

## 2 Results

### 2.1 Experimental design

Human observers (N = 10) performed a forced-choice orientation-discrimination task on a briefly-displayed peripheral grating (Fig. 1C). Observers fixated a central cross on a screen, while eye position and pupil area were measured. On easy trials, the grating was tilted far from vertical and observers achieved ∼93% accuracy on average. On hard trials, the grating’s tilt was close to vertical and observers achieved ∼70% performance on average. The task difficulty was changed in alternate runs (75 trials/run, 4 s/trial). In a separate version of the task, trial duration was shortened to 2 s/trial. We fit a computational model (Fig. 1B) to measurements of pupil size and saccades (including microsaccades), generating estimates of arousal level.

A correct estimate of arousal should be invariant to trial duration, but vary with task difficulty and behavioral performance. Manipulating task difficulty should alter arousal, as one needs to be more alert to achieve good performance on a more difficult task. Small changes in trial duration (i.e., 2 vs. 4 s), however, should not substantially alter arousal.

### 2.2 Saccade rate predicted the dynamics of the task-evoked pupil response

Saccade rate, transformed using our linking model, accurately predicted the task-evoked pupil response for different tasks and observers. The probability of saccade occurrence changed over a trial (Fig. 2A, Fig. S2). It fell after trial onset and rose again thereafter. We assumed that a saccade occurred each time a noisy “generator function” crossed a fixed threshold (Fig. 2B; *Methods*, Eqs. 1-2). Assuming that the variability in the generator function across trials was standard normal (i.e., i.i.d., additive Gaussian noise with standard deviation *σ* equal to one), we solved for its expected value across trials (Fig. 2C, see *Methods*, Eq. 6). That is, the expected generator function was simply a non-linearly compressed version of saccade rate, where the non-linearity was an inverse cumulative Gaussian. Next, the expected generator function was mean-subtracted, amplified by a gain, and low-pass filtered using a linear filter (Fig. 2D, black dashed curve; *Methods*, Eqs. 3, 7), yielding a prediction of the trial-averaged pupil response (Fig. 2E, yellow curve). We estimated gain as the scale factor that resulted in the best match (least squares, i.e, linear regression) to the pupil data. To estimate the linear filter, we measured pupil size following each saccade (Fig. 2D, purple curve) and fit it with a parametric function (Fig. 2D, black curve; *Methods*, Eq. 8; Fig. S3A). An additive offset, the average pupil size over a run, was added to the prediction of pupil size. For each observer, we modelled the post-saccade refractory period by adjusting the saccade rate function (see *Methods*). The measured task-evoked pupil response was a close match to the model prediction for all observers (Fig. S4; median *R*^2^ across all runs and observers, 81%; mean *R*^2^, 71%; max *R*^2^, 98%; min *R*^2^, 0%; N = 143 runs; see *Supplementary Information* for comments on individual differences).

**Figure 2:**
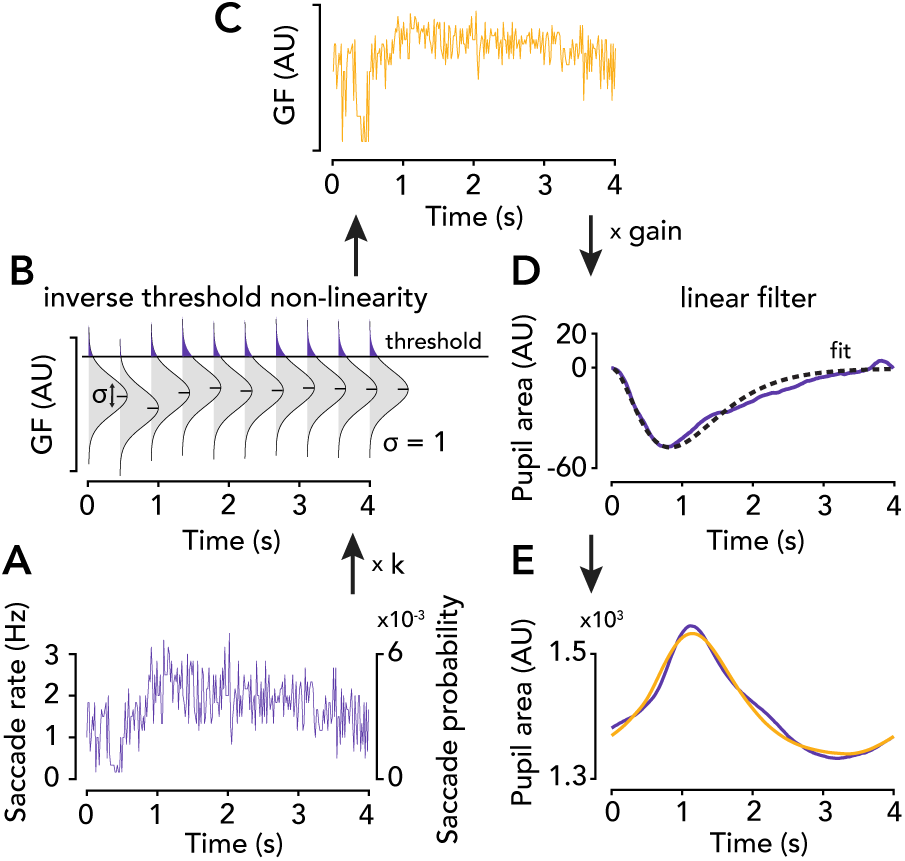
Model fitting. (**A**) Purple curve, saccade rate in Hz (left axis) or probability (right axis), averaged across trials, Gaussian-blurred for visualization. To model post-saccade refractory period, the measured saccade rate function is multiplied by *k*, the shape parameter of a gamma distribution fit to inter-saccade time intervals. (**B**) Inverting the threshold non-linearity. Grey distributions, standard normal noise (across trials) corrupting generator function (“GF”). Long horizontal black line, threshold. Purple tail, integral equal to the mean saccade rate in panel A. Short horizontal black lines, expected value of the generator function, 10 samples only (plotted in full in panel C). (**C**) Estimate of expected generator function. Gaussian-blurred for visualization. Mean-subtracted and multiplied by gain before filtering. (**D**) Linear filter. Black dashed curve, estimated linear filter — a gamma-Erlang function fit to the saccade-locked pupil response (purple). (**E**) Purple curve, task-evoked pupil response. Yellow curve, model prediction. Gain is the scale factor that gives the closest match between the data and model prediction.

### 2.3 Gain and arousal

#### 2.3.1 Gain, but not offset, scaled with task difficulty

We used our linking model to make predictions of the task-evoked pupil response separately for easy and hard trials. Accuracy in the orientation discrimination task was much higher for easy than hard runs, confirming that our manipulation of task difficulty was effective (t-test: t = 39.75, p = 2.71e-49, N runs = 143). The model predictions and data were consistent in shape and amplitude for both easy and hard trials (Fig. 3B, Fig. S4). Indeed, the model’s goodness of fit was indistinguishable for easy and hard runs (t-test: t = -1.14, p = 0.26).

**Figure 3:**
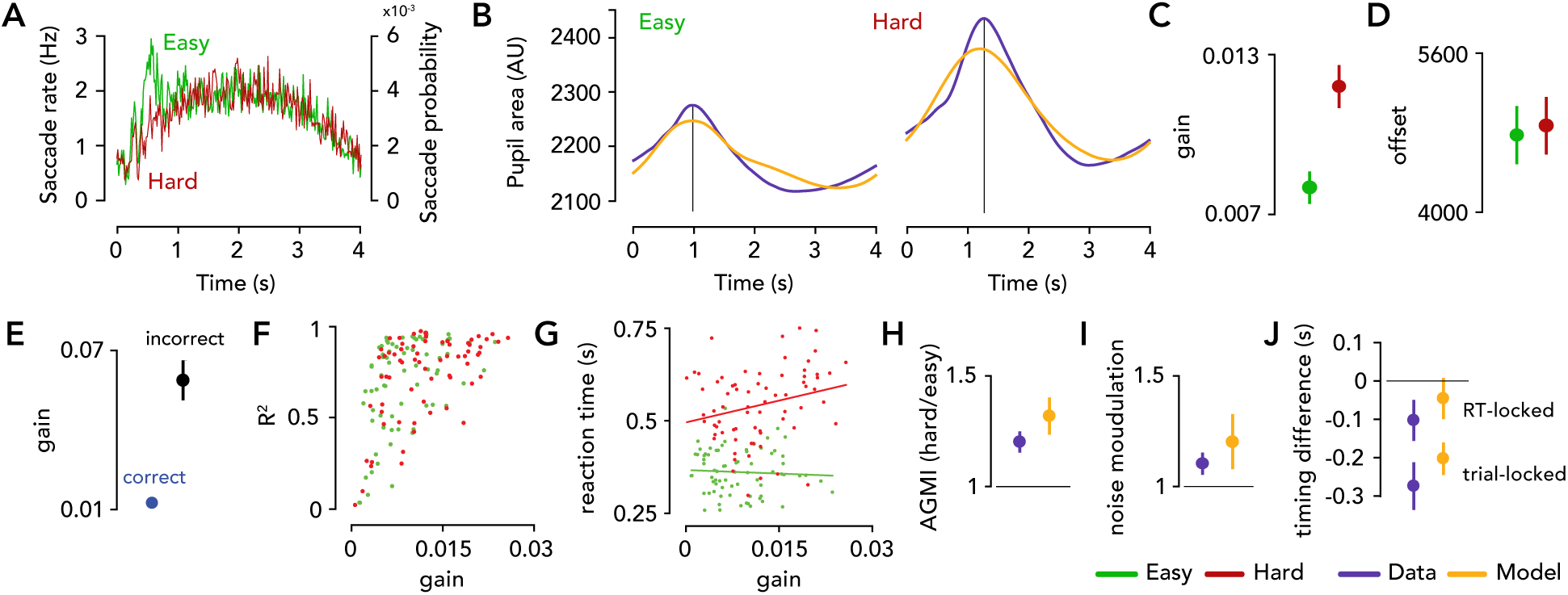
Model predicts difficulty-dependent modulations in the amplitude, timing, and variability of the task-evoked pupil response. (**A**) Saccade rate was similar for easy and hard trials, except immediately after trial onset. Curves, saccade rate in Hz (left axis) or probability (right axis), average over observers. Green, easy runs. Red, hard runs (same color convention throughout). (**B**) Task-evoked pupil response was larger in amplitude and more delayed when the task was harder. Purple curve, task-evoked pupil response. Yellow curve, model prediction. Left, easy. Right, hard. Vertical grey line, time-to-peak. (**C**) Gain scaled with task difficulty. Filled circles, mean gain across all runs and observers. Lines, two SEM (N = 143). (**D**) Offset did not vary with task difficulty; same format as C. (**E**) Gain was much higher for incorrect (N runs = 51) than correct trials (N = 115). (**F**) Model’s goodness of fit depended on gain (see *Supplementary Information*). *R*^2^ was variable for low gain and concentrated near 1 for high gain. Each circle, one run (N runs = 143). (**G**) Gain was correlated with reaction time and sign of correlation depended on difficulty. Lines, best-fit regression lines. (**H**) Model predicted amplitude modulation of the saccade-locked pupil response with task difficulty. Dots, amplitude or gain modulation index (AGMI), i.e., ratio (hard:easy) of minimum of saccade-locked pupil response or ratio (hard:easy) of best-fit gain. Errorbar, two SEM (N = 15). Purple, data. Yellow, model prediction. (**I**) Modulation of pupil noise with task difficulty. Dots, ratio (hard:easy) of pupil noise level or of best-fit gain. Errorbar, two SEM (N = 10). (**J**) Model predicted pupil response timing. Dots, mean difference (easy-hard) in the time-to-peak (s) of the pupil response. Errorbar, two SEM (N = 15). Top, reaction-time-locked pupil response. Bottom, trial-onset-locked pupil response.

Arousal should scale with task difficulty, i.e., a more difficult task requires greater alertness to achieve good performance. Gain, but not offset, was strongly modulated by task difficulty (Fig. 3C,D). We fit one free parameter per run, gain. The best-fit gain values, pooled across observers, were significantly larger for hard than easy runs of trials (t-test: t = 3.38, p = 9e-4, N = 143 runs). This difference remained significant in a logistic regression (F(141) = 10.27, p = 1.7e-3, N runs = 143), in which we accounted for systematic differences in gain across observers and experimental sessions (see *Supplementary Information, Statistical Analysis* for details). Offset, on the other hand, was not significantly modulated by task difficulty (logistic regression: F(141) = 0.05, p = 0.83, N = 143). Gain scaled with task difficulty, an expected outcome if gain estimates arousal.

#### 2.3.2 Gain was much higher on error trials

A correct estimate of arousal level should vary with behavioral performance, as arousal is known to influence behavior. We fit our linking model separately to pupil data from correct and incorrect trials (Fig. 3E, Fig. S5), expecting that gain would be higher on incorrect trials due to the tone feedback immediately following the button press, which was alarming to observers (particularly on easy runs). Because there were fewer incorrect than correct trials overall, and hence fewer saccades and pupil responses to constrain the model estimates, the fits were poorer overall on incorrect trials (median *R*^2^, 58.16% for incorrect trials; 80.74% for correct; mean *R*^2^: 55.93% and 70.87%). That said, there were many runs with good fits. We used only runs with an *R*^2^ over 50% (81.38% of runs for correct trials and 37.93% of runs for incorrect trials) for this analysis. Gain was 5.58x higher on incorrect than correct trials (t-test: t = 7.22, p = 1.59e-11, N runs = 118 correct, 55 incorrect; Fig. 3E) and this difference was evident separately in hard and easy runs (Fig. S5D,E). Gain modulation with behavioral accuracy was also evident in the trial-averaged pupil response, which had a larger amplitude on incorrect than correct trials (Fig. S5A). On the other hand, saccade rate was indistinguishable for correct and incorrect trials (Fig. S5C), on average across observers (cluster based permutation test: p > 0.05 for all time points). Likewise, the generator function was similar, particularly before the behavioral report occurred (Fig. S5B).

#### 2.3.3 Gain, but not offset, varied with reaction time

Gain, but not offset, was modulated by reaction time (average across trials in a run) and the direction of modulation was reversed for easy and hard runs (Fig. 3G). That is, the interaction of difficulty and gain was a significant predictor of reaction time (gain x difficulty: F(139) = 5.80, p = 0.017, N = 143; regression analysis; see *Supplementary Information*). On easy runs, observers often reported the task was trivial such that they became sleepy or distracted, and reaction time was faster when gain was higher and slower when it was lower. The latter may reflect a state of disengagement. And vice versa, on hard runs, observers were slower when gain was higher and faster when it was lower. This may reflect increased cognitive load (and thus, deliberation time) on trials when orientation was close to vertical. The three-way interaction between difficulty, gain, and accuracy was also a significant predictor of reaction time (regression: F(135) = 6.49, p = 0.012, N = 143). This corresponded with the observation that on the easy runs on which observers had the worst accuracy, the negative relationship between gain and reaction time became even more pronounced (than on the runs with higher accuracy). We repeated these regressions but predicting offset instead of gain, and all predictions were non-significant (offset x difficulty: F(139) = 0.06, p = 0.80091; offset x difficulty x accuracy: F(135) = 0.52, p = 0.47; N = 143). Reaction time depended on gain, as expected if gain estimates arousal level.

#### 2.3.4 An independent prediction: amplitude modulation of the saccade-locked pupil response with task difficulty matches modulation of gain

Our model predicts that the amplitude of the saccade-locked pupil response scales with gain (Fig. S6B,D). We measured this prediction (Fig. 3H) as the ratio (model:data) of the ratio (hard:easy) of the minimum of the saccade-locked pupil response (Fig. S3B) and the ratio (hard:easy) of the mean best-fit gain. A perfect model prediction would yield a ratio (model:data) of 1. The median ratio across observers was 1.13 (mean, 1.11; SEM, 0.09; for observers O1-O10: 1.06, 0.87, 1.188, 1.15, 1.13, 0.82, 0.78, 1.14, 1.22, 1.73). We estimated the linear filter used in our model fits from the run-averaged saccade-locked pupil response, throwing away its run-to-run variability. Therefore, this is an independent prediction of the model — that is, a prediction about data that was not used to fit the model parameters.

#### 2.3.5 An independent prediction: pupil noise scaled with gain

In our model, trial-to-trial variability in pupil size (“pupil noise”) is inherited from noise in the generator function. We assume that this input noise is additive and Gaussian, such that when it is amplified by a gain *g*, its new standard deviation is equal to *g* (see *Methods*, Eq. 5). The output of any linear filter acting on a Gaussian input will also be Gaussian [77], with the property that if you double the input noise, the output noise will also double (i.e., a linear mapping). Thus, our model predicts that if the best-fit gain is x times larger for hard than easy trials, then the standard deviation of pupil noise should also be x times larger. Note that this is an independent prediction of the model, as we estimated gain based only on the trial-averaged pupil response, ignoring its trial-to-trial variability.

For each observer, we quantified the level of pupil noise by aligning the pupil responses from each trial and computing the standard deviation of pupil size across trials for every time point from 0 to 4 s (Fig. S7A; see *Supplementary Information, Pupil noise analysis and simulation* for details). The pupil noise level was significantly higher for hard than easy runs for 7/10 observers (paired t-test, p < 5e-4, N observers = 10). For one observer, O5, the pupil noise level was lower for hard than easy runs, but this matched their best-fit gain, which was also smaller for hard than easy. The distribution of pupil noise closely resembled a Gaussian (Fig. S7B,C), validating our noise model and use of standard deviation as a measure of the noise level. The disparity in the pupil noise level between easy and hard runs (“noise modulation”) was approximated well by a single scale factor, matching our model’s assumption of stationary noise within a trial (Fig. S7A).

To quantify our model’s ability to predict modulation of pupil noise with task difficulty, we computed three ratios for each observer: (1) empirical noise modulation, the ratio (hard:easy) of the average pupil noise level (across time), (2) predicted noise modulation, the ratio (hard:easy) of the best-fit gain (Fig. 3I), (3) the ratio between empirical and predicted noise modulation. If ratio #3 is 1, it means that our model predicted pupil noise modulation perfectly. The median ratio #3 (data:model) across observers was 0.95 (mean = 0.89, SEM = 0.09, ratios for each observer O1-10: 0.66, 1.09, 0.64, 1.29, 1.06, 1.19, 1.15, 0.84, 0.55, 0.46). Although noise modulation was slightly larger in the model than data (Fig. 3I, Fig. S7A), the predictions were largely in the right direction and errors were small.

### 2.4 Timing and the generator function

#### 2.4.1 Dynamics of saccade-locked pupil response were modulated by trial duration

The model predicts that the saccade-locked pupil response is different (distorted) from the underlying linear filter because of the threshold nonlinearity for generating saccades. In the data, we found that the nature of this distortion depended on trial duration (i.e., via differences in the generator function), and was particularly bizarre for the 2 s task (Fig. S6C), where it caused rippling. Our model closely reproduced this task-dependent distortion (Fig. S6B,D, Fig. S3), perhaps the most idiosyncratic feature of the data, providing a strong confirmation that the model architecture is correct.

#### 2.4.2 Generator function dynamics were modulated by the timing of the task and behavioral response, but largely invariant to task difficulty

We hypothesize that the generator function reflects the timing of the task and behavioral response, but is invariant to arousal level. If so, we should find that the generator function changes with trial duration and reaction time, but is the same regardless of task difficulty.

We quantified differences in the generator function between easy and hard trials (Fig. S8) by computing the separation between the generator functions for easy vs. hard trials at every time point from 0 to 4 s (mean generator function was first averaged over all easy or hard runs, mean-subtracted, and Gaussian-blurred over time). We chose d’ as a measure of distribution separation, which in this case (standard deviation 1; Eq. 1) was simply equal to the difference between the two generator functions. We observed a spike in the generator function on easy but not hard trials at around 0.5 s after trial onset, for most observers. We computed the time of maximal separation as the maximum d’ (over time). Indeed, O1, O2, O3, O4, O5, O7, and O9 had a large increase in d’ (> 1.43) sometime between 0.40 and 0.65 s. For context, the average d’ over the full trial ranged from 0.30 to 0.47 across observers.

To test whether these differences were due to differences in reaction time between easy and hard trials (Fig. 4A,B), rather than difficulty per se, we conditioned the generator function on reaction time, task difficulty, or both (Fig. 4A). Data were pooled across observers (N = 10). The initial spike in the generator function was larger when reaction time was faster, for both easy and hard trials (Fig. 4A). Indeed, there was a region of maximal separation (d’) between the generator functions at around 250 ms for both easy and hard trials. We next compared the generator functions (easy vs. hard) for fast (Fig. 4C) or slow (Fig. 4D) reaction times separately, and found smaller differences. This suggests that “difficulty-dependent” modulations in the generator function (Fig. S8) were actually driven by differences in reaction time between easy and hard trials. Therefore, the generator function is modulated primarily by reaction time, rather than task difficulty.

**Figure 4:**
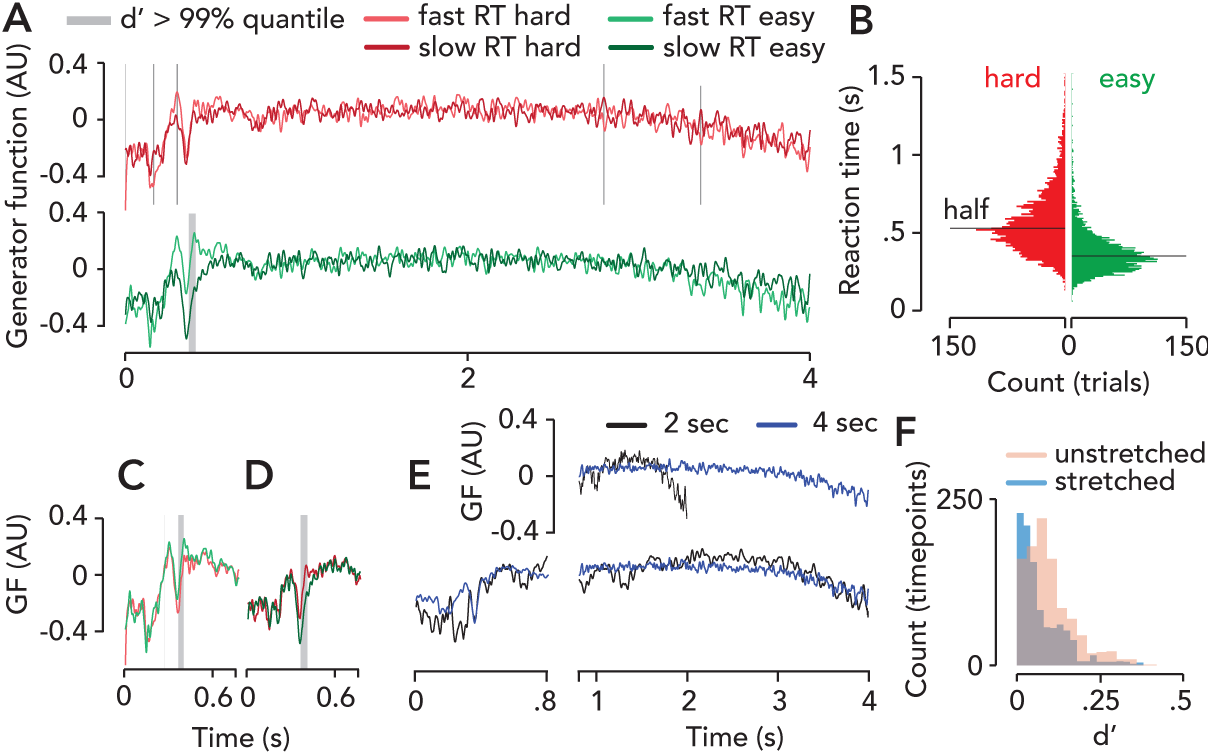
Generator function was modulated by timing of the task and behavioral response. (**A**) Generator function (“GF”) for fast and slow reaction times (median-split). GF computed from average saccade rate (N observers = 10, N trials = 3450). Red, hard runs. Green, easy runs. Light colors, fast RT (below median RT). Dark colors, slow RT (above median RT). Grey regions, times of maximum separation between GFs (i.e., > 99% quantile of d’ over time). (**B**) Distribution of reaction times for easy and hard trials (N observers = 10, N trials = 3450). Horizontal black lines, medians. (**C**) GF for fast reaction times (hard vs. easy trials), re-plotted from A. There were no large separations beyond 0.8 s, so only first 0.8 s shown. (**D**) Same as C, but for slow reaction times. Difficulty-dependent modulations in GF were smaller than reaction-time-dependent modulations (compare differences between curves in panels A and C). (**E**) GF for 2 or 4 s task (N observers = 10; N runs = 97 for 4 s, 39 for 2 s). Black curve, 2 s; blue curve, 4 s. First 0.8 s are in original time base. Top, unstretched data (i.e., raw data). Bottom, from 0.8 to 4 s, data from 2 s task is time-stretched by a factor of two, revealing that the GF is similar for 2 s and 4 s tasks after time-stretching. (**F**) Distributions of d’ between the GFs corresponding to the 2 s and 4 s tasks, for 0 to 2 s after trial onset (N = 1000 timepoints). Cyan, d’ when GF for 2 s task was time-stretched beyond 0.8 s after trial onset. Pink, d’ for raw data (i.e., unstretched).

The generator function was also modulated by the timing of the task, stretching with trial duration. Five new observers performed the task with 2 or 4 s long trials, in separate blocks. We compared the generator functions between the 2 and 4 s tasks (Fig. 4E, Fig. S11C), with and without time-stretching (Fig. 4E,F). Data were averaged across observers (N observers = 10, N runs = 97 for 4 s task, 39 for 2 s task). Specifically, we time-stretched the generator function (Fig. 4E) from the 2 s task by a factor of two for all time points beyond 0.8 seconds (i.e., after nearly all button presses had occurred, Fig. 4B, and when the observer was expecting next trial to begin). Then we computed the distribution of d’ across the first 2 seconds of the trial, between the 2 and 4 s tasks, before and after time-stretching (Fig. 4F). D’ was significantly lower for the time-stretched generator function (t-test: t = 10.42, p = 3.29e-24) and this comparison was still significant after log-transforming the skewed distributions to make them more Gaussian (t = 10.32, p = 8.72e-24). There were also differences in the generator functions between the 2 and 4 s tasks before 0.8 seconds (i.e., the never-stretched region, during which stimulus onset, button press, and feedback occur); the maximum d’ of 0.38 occurred 290 ms after trial onset (Fig. 4E).

#### 2.4.3 Pupil response timing was predicted by reaction-time-dependent modulations in the generator function

When present, the transient spike in the generator function on easy trials (Fig. S8) correctly predicted the advanced time-to-peak of the task-evoked pupil response (Fig. 3B). The task-evoked pupil response peaked earlier for easy than hard trials (Fig. 3B, Fig. S4). We hypothesize that this timing difference is driven by differences in the distribution of reaction times (i.e., advanced timing and lower dispersion for easy trials), which is reflected in the generator function’s dynamics (i.e., an early spike in easy but not hard trials). To test this, for each observer we (1) computed the pupil response time-locked to trial onset (“trial-onset-locked”, equivalent to “task-evoked”) or to button press (“RT-locked”) (Fig. S9B,D), computed the difference (easy minus hard) in the response time-to-peak, (2) estimated the generator function time-locked to trial onset or reaction time (Fig. S9A), used our model to predict pupil size (Fig. S9C,E), and computed the difference (easy minus hard) in the model response’s time-to-peak. If our explanation is correct, the model prediction should capture the timing differences (easy vs. hard) seen in the data.

For the trial-onset-locked pupil response, the timing difference (time-to-peak for easy minus hard) was statistically indistinguishable for model and data (t-test: t = -1.09, p = 0.30, N = 15) with an average of -274.3 ms for data (SEM, 62.3) and -202.7 ms for model (SEM, 42.4), i.e., only a 71.6 ms difference between model and data (Fig. 3J). After removing one outlier (O7, who had an abnormal biphasic pupil response, causing the time-to-peak to be very low), this difference dropped to 25 ms. For the RT-locked pupil response, the timing difference was also statistically indistinguishable between model and data (t-test: t = -0.81, p = 0.43, N = 15), with an average of -102.8 ms for data (SEM, 53.6) and -46.7 ms for model (SEM, 55.4), i.e., a 56.1 ms difference between model and data. After removing the outlier (O7), this difference dropped to 10.1 ms. Thus, timing differences in the task-evoked pupil response were driven by a spike in the generator function present on easy but not hard trials, which was linked to faster reaction times. When the spike was also present on hard trials (i.e., in the RT-locked generator function), these timing differences disappeared. This demonstrates that pupil response timing reflects reaction-time dependent modulations in the generator function.

### 2.5 Model comparison

#### 2.5.1 Pupil response amplitude depends jointly on gain and generator function

The amplitude of the task-evoked pupil response, commonly used as a heuristic measure of arousal level, depends on both gain and the dynamics of the generator function. We separately quantified the components of pupil response amplitude (i.e., maximum pupil size relative to trial onset baseline) due to the gain and generator function by analyzing our model’s prediction before and after gain modulation for easy versus hard runs (Fig. S10). The pre-gain model prediction captured some of the differences in pupil response amplitude between easy and hard runs, both for individual observers (Fig. S10A) and on average over all observers (Fig. S10B). But the difficulty-dependent change in the generator function and the difficulty-dependent change in gain were *both* needed to account for the pupil size measurements.

#### 2.5.2 Comparison with existing models

Our model generalized between different tasks, whereas the consensus model could not. To assess generalization performance, we had a group of observers (N = 5) perform two versions of the task which differed only in trial duration (2 or 4 s). Pupil and saccade data were markedly different for the two trial durations (Fig. S11A,B), but behavioral accuracy was nearly identical (mean accuracy across runs, 2 vs. 4 s task, hard: 72.93% vs. 72.46%; easy: 96.67% vs. 98.78%). A correct model linking arousal and pupil size should recover a similar estimate of arousal (i.e., similar parameter estimates) for both tasks, despite the different pupil size dynamics. Therefore, we tested if the parameters estimated from one data set applied to the other nearly as well, generating accurate predictions of pupil size. To do so, we fit either our model or the consensus model to the 2 s data and evaluated the best fit parameters on the 4 s data, and vice versa. This was done separately for easy and hard runs, for each observer. We quantified generalization performance as the median reduction in *R*^2^ between the out-of-sample prediction vs. in-sample fit. For our model, we used the average gain across all easy or hard runs for the out-of-sample prediction. For the consensus model, we fit the amplitudes of “arousal-related” impulses at trial onset and button press and of a “time-on-task” boxcar (Fig. S11D). We used these three amplitude parameters (averaged over easy or hard runs) for the out-of-sample prediction. For our model, the median reduction in *R*^2^ across observers was 14.67% (reductions for each observer, 4->2 s prediction: 10.06, 17.21, 0.60, 36.42, 19.94%; 2->4 s: 12.14, 20.96, 5.40, 7.01, 124.66%). For the consensus model, the median reduction in *R*^2^ across observers was 46.89% (reductions for each observer, 4->2 s prediction: 47.07, 65.65, 62.33, 16.94, 59.49%; 2->4 s, 21.645, -16.91, 117.31, 7.49, 46.73%). For our model, the median *R*^2^ values for the in-sample fits were 86.27% (2 s) and 78.62% (4 s). For the consensus model, they were 94.16% (2 s) and 39.14% (4 s). Taken together, these results suggests that the consensus model overfits and that its predictions are not self-consistent.

The primary reason that the consensus model fit the 2 s data better than the 4 s data was the presence of a large pupil constriction relative to pre-trial baseline around 1 s after trial onset in the 4 s, but not 2 s task (Fig. S11A) [40]. The consensus filter is dilatory and so can not capture such constrictions at all (Fig. S11D). On the other hand, our model predicted this large constriction on the 4 s data, as well as the much smaller constriction in the 2 s data (Fig. S11A). The reason is simply the prolonged generator function (Fig. S11C), and hence prolonged input to the constrictory linear filter. It’s clear a priori that adding a second linear system with a constrictory filter, as some have proposed [40, 42], would make the consensus model’s in-sample fit in the 4 s task much better, but it would also make the model (even) less generalizable (as there would be a near-zero amplitude best-fit “button press” impulse for the 2 s task and a larger, negative amplitude impulse for the 4 s task).

#### 2.5.3 Pupil size is a linear transformation of the generator function, not trial events

Model comparisons were performed to assess whether the pupil is a shift-invariant linear transformation of impulses time-locked to trial events, as existing models assume, or of the gain-modulated generator function, as our model assumes. To test these assumptions, we computed the ratio of the best-fit scale parameters (i.e., gain or impulse/boxcar amplitudes) from the 2 and 4 s tasks. We refer to this ratio as the additivity/shift-invariance or “ASI” index. If pupil size is a linear shift-invariant transformation of its input, this ratio should be 1. For the consensus model, the mean ASI index across observers was 2.45e+11 (SEM, 1.60e+11) for the boxcar, 1.54e+10 (SEM, 1.54e+10) for the trial onset impulse, and 0.61 (SEM, 0.13) for the button press impulse. The medians were 0.53, 0.57, and 0.58, respectively. The reason for the large mean values is that the amplitude estimates were unstable because the three parameters traded off with each other. For our model, the mean ASI index across observers was 0.9323 (median, 0.80; SEM, 0.12) and was statistically indistinguishable from 1 (t-test: t = -0.1720, p = 0.8673, N = 5). Next, we computed a homogeneity/shift-invariance or “HSI” index. This was the ratio (4 vs. 2 s) of the quotient of the hard and easy best-fit gain values (our model) or amplitudes (consensus model). If pupil size is a linear shift-invariant transformation for its input, the HSI index should be 1. For the consensus model, the mean HSI was 0.09 (SEM, 0.06) for the boxcar, 1.30e+11 (SEM, 1.30e+11) for the trial onset impulse, and 6.06e+10 (SEM, 6.06e+10) for the feedback impulse. The medians were 0.01, 3.56, and 2.16, respectively. For our model, the mean HSI index was 1.13 (median, 1.17; SEM, 0.18) and was statistically indistinguishable from 1 (t = 0.30, p = 0.78, N = 5). Taken together, these results suggest that the pupil is a shift-invariant linear transformation of the gain-modulated generator function, not trial events. These results also show that gain was stable across tasks, suggesting that gain reflects arousal level, but not task structure.

## 3 Discussion

We propose a model of the task-evoked pupil response, which capitalizes on the hypothesis that a common input drives both saccades and pupil responses, i.e., the “common drive hypothesis” [46, 10, 45, 47, 26, 21]. Our model estimates two main components, the generator function (i.e., the common input) and gain. We hypothesize that the generator function reflects task timing and gain reflects arousal level. To test this idea, we fit our model with saccade and pupil size measurements from a visual orientation-discrimination task in which task difficulty and trial duration were varied. The model accurately predicted pupil size from saccade rate. Estimates of gain were modulated by task difficulty and behavioral accuracy, but not by trial duration. On the other hand, the estimated generator function was modulated by the trial duration, but not by difficulty or accuracy. Both the gain and generator function were modulated by reaction time. Existing models assume that pupil size is a linear transformation of arousal-related impulses at the time of trial events. Such models failed to generalize across tasks with different trial timing despite achieving nearly perfect in-sample fits, demonstrating that this assumption is false and that these models are over-fitting. On the other hand, our model achieved good generalization and fits. One implication is that it is necessary to measure both saccades and pupil size to correctly estimate the input to the pupil and to thereby constrain estimation of arousal. Our model provides a unified explanation for a number of seemingly disparate observations, including entrainment of pupil size and saccade rate to task timing, amplitude modulation of pupil responses with task difficulty, modulation of trial-to-trial pupil noise with task difficulty, a constrictory pupil response time-locked to saccades, task-dependent distortion of this saccade-locked pupil response, and reaction-time dependent modulations of pupil response timing. These results provide strong support for the hypothesis that a common input (the generator function) drives saccades and pupil size, a saccade occurs when the generator function crosses a threshold, the task-evoked pupil response is a gain-modulated and low-pass filtered transformation of the generator function, and that arousal corresponds to the gain.

Over the past 80 years, measuring pupil size has become a popular tool for inferring arousal level [21]. Inexpensive, non-invasive, and high fidelity, pupil size recordings have many advantages over the alternatives, i.e., measurement of spiking activity in neuromodulatory loci [26, 14], whole brain imaging [32, 78], and in-vivo biochemical recordings [79, 80]. Indeed, a large number of recent publications have linked pupil size with behavior [41, 11, 34, 81, 82, 83, 40, 32], or neural signals [29, 26, 10, 25, 31, 30, 32]. Approaches to estimating arousal from pupil size fall into two categories: heuristic and model-based. A popular heuristic analysis in systems neuroscience involves computing the correlation (or regression) between the time course of pupil size and neural activity in some population of interest, where pupil size is treated as a measurement of arousal level [32, 29]. The implicit assumption is that the neural input to the pupil tracks with arousal level and that delays and temporal summation caused by the sluggish response of the iris muscles [84, 85] don’t substantially obscure the underlying correlation between the neural signals. Another popular heuristic approach estimates arousal level based on the height of the pupil response (i.e., max or max-min size) [86, 33].

The most popular model-based approach is a linear systems model, which assumes that the pupil response is a low-pass-filtered transformation of arousal-related neural activity aligned to task events [34, 35, 50, 87, 36, 37, 39, 38, 88, 42]. This assumption is often justified based on the physiological findings that locus coeruleus (LC) neurons spike at the time of salient events [14, 27] and that LC activity or microstimulation precedes pupil dilation [25, 89, 26]. Some highly-cited papers thus claim it is possible to infer activity in the LC-NE system from pupil size [90, 91, 76, 92]. This consensus model is nearly identical to one that has been used in functional magnetic resonance imaging (fMRI), in which the strength of neural activity is inferred by regressing a prediction, the convolution of the hemodynamic response function (HRF) with impulses at trial event times, on the fMRI signal. In fMRI, this approach was validated by the observation that the fMRI signal approximates a shift-invariant linear transformation of local spiking activity [93] and that neural activity is time-locked to stimulus onsets in many sensory areas of the brain. But linearity of pupil size with respect to its purported input has never before been tested. Furthermore, it is unclear whether the neural input to the pupil is aligned with trial events. Our model comparisons suggest that both assumptions are false — the consensus model’s parameters did not generalize across tasks with different trial timing, demonstrating that pupil size is not a linear transformation of an input time-locked to trial events. Earlier versions of the consensus model [43, 44] assumed that the input to the pupil were impulses that could occur at any time (analogous to our generator function) rather than being time-locked to task events. We found, however, that the parameters of such models were not recoverable (see *Supplementary Information, Parameter recovery: deconvolution method of Wierda and colleagues*), suggesting that these models are underconstrained for pupil size measurements alone.

The challenge of estimating arousal from the task-evoked pupil response is that the input to pupil and arousal level are both unknown, making the problem under-constrained. The consensus model constrains the problem by assuming that the input’s timing is equal to the task timing (which is known because it is set by the experimenter) and that the input’s form is equal to some particular parametric form (e.g., impulse, boxcar). We adopted a different approach, in which we used a model to estimate the input to the pupil from measurements of saccades rather than assuming its timing and form. By analogy, fMRI measurements in early visual cortex are analyzed by adopting a model that maps from the pixel intensities of an image to the evoked neural activity [94]; such models are based on the known anatomy and physiology of the brain and include multiple stages comprising non-linear and linear operations [95]. The advantage of such “image-computable” models is the ability to make predictions of a measurable signal (i.e., the fMRI signal) from another measurable physical signal (i.e., an image) and to compare the prediction to data under many conditions, putting a constraint on the possible operations in-between. Here, we took a step toward adopting an analogous theoretical framework for estimating arousal from pupil size. Future work is needed to develop a model that links task structure to the generator function. Currently, this mapping is unknown. If this can be achieved, then it will be possible to combine our model with this one, yielding a model that predicts pupil size from the timing of task events, i.e., a “task-computable model.”

Our results demonstrate that the amplitude of the task-evoked pupil response depends on both the generator function and gain (Fig. S10), implying that the same measured pupil response can be due to many different combinations of the two components. Hence, we used model fitting and comparisons to tease out whether pupil size modulation was due primarily to changes in gain, changes in the generator function (i.e., the input to the pupil), or to both. These results prompt a re-interpretation of many previous findings relating the amplitude of the task-evoked pupil response to various psychological processes, including memory, learning, cognition, attention, and decision-making. Our model makes a strong prediction that can aid in this re-interpretation: psychological processes that influence both saccade rate and pupil size do so via the generator function. Processes that influence pupil size, but not saccade rate, do so via gain [61]. Some processes will influence both. Our model is needed to quantify these contributions.

We speculate that the generator function, the driver of saccades and pupil size, is influenced by multiple aspects of cognition. We manipulated trial duration in our study and found that during the inter-stimulus interval, the generator function stretched with trial duration (Fig. 4E), whereas between the stimulus onset and behavioral response, it was modulated in a more complex way (Fig. 4E, Fig. S11C). It is well known that trial duration influences task engagement and anticipation [96, 97, 98], as well as uncertainty, if duration is variable [99]. Thus, modulations in the generator function may reveal the extent of task engagement and anticipation. In our model, the generator function is simply a transformed version of the trial-averaged (micro)saccade rate. Thus, previous results revealing that (micro)saccade rate is modulated by temporal expectation and attention [73, 60] also support this conclusion. Extending this logic, it is well accepted that many cognitive processes affect (micro)saccade rate, including language processing [100], sound categorization [101], working memory [75], emotional responses [102], and voluntary saccade planning [103, 104]. Thus, although we only tested the effect of trial duration on the generator function in this study, we expect that the generator function reflects many other aspects of cognition.

### 3.1 Physiological basis

We can only speculate about the physiological basis of the four main components of our model: the generator function, threshold, gain, and linear filter. We offer a disclaimer that our model is simple and abstract by design. Multiple model components could correspond to one part of the neural circuitry, or vice versa; one model component could correspond to multiple parts of the circuitry. Further anatomical, physiological, and psychophysical experiments are needed to determine these correspondences.

We speculate that the generator function corresponds with the pooled activity of superior colliculus neurons. Specifically, we hypothesize that: (1) SC drives both saccades and pupil responses; and (2) SC encodes a cognitive representation of the task, including its timing. Support for the first of these hypotheses is evidence that the intermediate/deep layers of the superior colliculus (SCi) are a common input to the muscles that control pupil responses and saccades [46, 10, 26, 45, 21]. SC projects directly to the preganglionic Edinger-Westphal (EW) nucelus, which controls pupil constriction and is most widely known for its role in mediating both the pupil’s accommodative near response and light reflex [77, 84, 105, 10]. SC, most known for its role in saccade generation [106, 107], is involved more generally in orienting behaviors (including eye, head and body movements) that guide target selection and spatial decisions [108]. But spiking in SC neurons is also locked to a pupil response [46, 26, 10, 62] and crucially, microstimulation of SC neurons (above or below the threshold for saccade generation) evokes a pupil response [46, 26, 45].

Support for the second of the above two hypotheses is evidence that SC neurons encode the saliency of visual, auditory and somatosensory events [109], thus allowing it to build a timeline of a task (i.e., reflecting task timing). Indeed, SCi neurons respond to behaviorally-relevant stimuli in a manner that is substantially invariant to their particular retinotopic location [110]. Optogenetically inhibiting SC, even ipsilaterally to a visual stimulus, causes increases psychophysical thresholds in a detection task, and increases in lapse and guess rates, suggesting that SC activity is important for maintaining task engagement [111]. Muscimol-mediated inactivation of SCi neurons causes biases in target selection, for saccadic, arm reach, and button press responses, despite not causing generic impairment of movement, demonstrating that SCi encodes target priority in an effector-independent manner [112, 113]. We speculate that a cognitive representation of task structure generated in higher cognitive areas, including dorsolateral prefrontal cortex (dlPFC) and anterior cingulate cortex (ACC), modulates superior colliculus activity, which in turn controls pupil size. This is consistent with previous proposals [10, 21, 54].

We speculate that the threshold in our model corresponds to the aggregate behavior of the circuit connecting SC, omnipause neurons, the brainstem burst generator, and the extraocular muscles. This circuit is known to initiate saccades and maintain eye position in the orbit [114]. The threshold in our model may approximate this entire circuit’s input-output relation — i.e., an abstraction of SC-driven saccade initiation. Thresholding of neural activity in SC or frontal eye field (FEF) is likely the most influential model of saccade initiation [115, 116, 117].

We speculate that the gain in our model corresponds with neuromodulation of activity in the preganglionic Edinger-Westphal nucelus (EW). A possible circuit mediating gain modulation is the direct inhibitory noradrenergic projection from LC onto EW [118, 2, 10], discovered in cat. However, the existence of this connection in humans is controversial [119]. There are also direct projections from LC onto SCi, which then in turn projects to EW both directly and indirectly (via the mesencephalic cuneiform nucleus) [10, 21]. The picture is enriched by the fact that cat EW in fact receives projections from many other neuromodulatory nuclei, including the ventral tegmental area, the pars reticulata of the substantia nigra, and the raphe nuclei [118, 120]. If these same afferent connections exist in humans, gain may be controlled by a number of neuromodulators, not just by the LC-NE system. This would explain why pupil responses have been observed locked to microstimulation of neurons in noradrenergic, serotonergic, and cholinergic brainstem loci [26, 20, 25, 54]. One could test this idea by measuring EW activity and pupil size with pharmacological manipulations that selectively affect these neuromodulators. If drugs that selectively affect the dopamine system, for example, gain-modulate EW activity, this would provide evidence that the pupil’s gain can be influenced by a number of neuromodulators, which collectively define “arousal level” [54]. This would also explain why pupil responses are modulated by reward and reinforcement learning [40, 121], which are commonly associated with the dopaminergic system.

We speculate that the linear filter in our model corresponds with the activity of the iris constrictor muscle, driven by the parasympathetic nervous system. In the parasympathetic pathway, the constrictor muscle is driven by cholinergic neurons arising in the ciliary ganglion, which are driven solely by afferents from EW [122, 123]. In the sympathetic pathway, the iris dilator muscle is under the control of adrenergic neurons in the superior cervical ganglion, which is driven by the intermediolateral nucleus (IML) of the spinal cord [124, 125, 21].

Pharmacological studies reveal that pupil responses during task performance and rest are primarily under parasympathetic control and that arousal modulates this circuitry. The task-evoked pupil response is nearly eliminated by a local parasympathetic antagonist (i.e., applied to the eye), but unaffected by a local sympathetic agonist [126], demonstrating that the task-evoked pupil response is primarily under parasympathetic control. Likewise, a *local* sympathetic antagonist or agonist has no effect on pupil size fluctuations at rest, while a local parasympathetic antagonist attenuates these fluctuations [6, 7]. Enriching this account, an orally-administered central alpha-2 agonist (i.e., decreases central NE) decreases pupil size fluctuations (at rest) in the light, but has no effect on pupil size in the dark. By contrast, an alpha-2 antagonist (i.e., increases central NE) increases pupil size fluctuations in the light, and has no effect in the dark [56, 57]. Together, this suggests that global neuromodulation (e.g., related to function of LC) can interact with the pupillary light reflex, which is primarily driven by the parasympathetic pathway [127, 128]. Supporting this idea, very high or low arousal attenuates the constrictory pupil response to a transient luminance change [1, p.480–481][127, 7, 129, 126, 58, 130, 18]. Physiological studies offer further support for the hypothesis that the task-evoked pupil response is primarily under parasympathetic control. In awake cat, a startling auditory tone leads to a characteristic rising-and-falling pupil response even if the sympathetic pathway is surgically removed, although its amplitude is attenuated by 1.5x and its dynamics are more low-pass compared to a normal pupil [55]. Stimulation of cortex or dorsal diencephalic regions (e.g., hypothalamus, one input to sympathetic pathway) of a sympathectamized animal causes inhibition of the pupil light reflex, whereas stimulation of superior cervical ganglion in an intact animal does not [127, 55]. Furthermore, in macaque, the modulatory influence of arousal level on the pupillary light reflex is nearly the same in a sympathectomized and intact eye [127]. These results support the hypothesis that inhibition of the parasympathetic pathway is the primary driver of the dilatory portion of the task-evoked pupil response [55]. One study of the light reflex found that simply passing the spike train of an EW neuron through a constrictory low-pass filter leads to an accurate prediction of pupil size [84], suggesting that the relation between EW activity and pupil size is linear. This supports our speculation about parasympathetic control of the linear filter in our model. Regardless of what drives activity in EW, whether its light, accommodation, sound, self-generated movement, or increased cognitive engagement, the pooled magnitude of EW activity (via ciliary ganglion) determines the total number of acetylcholine molecules released onto the iris constrictor muscles, and hence the magnitude of pupil constriction. Additional physiology experiments are needed that test linearity directly by simultaneously measuring pupil size and its neural inputs (i.e., the inputs to the dilator and constrictor muscles).

Our model assumes that pupil response variability is driven by the noise in the generator function and scales with gain. While previous studies have differentiated between variability in pupil responses at rest and during task performance [55, 1, 6], our model suggests that these share the same origin and are simply points along a continuum defined by the dynamics of the generator function. At one extreme (spontaneous), the generator function is flat in expectation, and so saccades and pupil size are unpredictable (Fig. S1A,B). This input may be systematic but endogenous (e.g., due to interoception or mind wandering) and therefore unpredictable to an outside observer, and cancels out when averaging over time. At the other extreme (task-evoked), the generator function entrains to the timing of external events (e.g., trial onsets) so saccades and pupil size modulations are more predictable (Fig. S1C,D,F). Importantly, pupil fluctuations are of a similar amplitude during task or rest (Fig. S1E). The assumption here is that the same (primarily parasympathetic) circuitry controls both task-evoked and spontaneous pupil fluctuations [126, 7]. Previous studies support this idea. Pupil noise in the two eyes is almost 100% correlated, implicating a noise source that drives both eyes, i.e., EW or higher in the circuitry [77]. Confirming this, pupil noise is attenuated if the nerve between the ciliary ganglion and its input, EW, is cut [1]. Fentanyl, an opioid agonist that attenuates inhibition of midbrain structures (like EW), depresses equiluminant pupil size fluctuations, suggesting that inhibition of EW produces these fluctuations [59]. This is consistent with a physiological study showing that EW has a high tonic firing rate, which is inhibited by a transient step-up in light level [84] — i.e., arousal may also inhibit tonic activity in EW. A statistical analysis by Stanten and Stark [77] revealed that pupil noise is multiplicative and Gaussian. This can arise from an additive Gaussian noise source followed by a gain and then subjected to a linear filter [131, 132, 77, 133]. Stanten and Stark found that the standard deviation of pupil noise scales with both light level and fixation depth (accommodation), suggesting that the same noise source is shared among all parasympathetic-mediated pupil responses. Consistent with this, we found that arousal modulated both gain and the standard deviation of pupil noise (Fig. S7), bolstering the idea that the task-evoked pupil response is parasympathetic-mediated and its noise is inherited from the gain-modulated generator function. Stark concluded that this noise source must be in the EW or higher [131, 132, 77, 133]. Our results demonstrate that variability in saccade generation and pupil responses is shared, i.e., we hypothesize that pupil noise must originate above EW, possibly in SC.

### 3.2 Possible confounds

The saccade-locked pupil response is not a luminance or accommodation response, nor an artifact of foreshortening caused by a change in eye position (a potential source of error in infared eye-trackers). Foreshortening error occurs when the detected pupil area or diameter is smaller (i.e., a squashed ellipse vs. a circle) due to gaze direction deviating from screen center (i.e., where the eye tracker is angled). The saccade-locked pupil response occurs with a delay of 0.9 s and takes 4 s to come back to baseline. An artifact based on eye position should arise and dissipate much more rapidly than 4 s. Moreover, many of the saccades measured in our study are corrective (i.e., they are toward the fixation point, correcting for drift). These small corrective saccades correspond to a relatively large *decrease* in pupil size, as opposed to the small increase predicted by a reduction in foreshortening. Furthermore, we found that the amplitude of the saccade-locked pupil response was not systematically related to saccade amplitude (see *Supplementary Information, Saccade amplitude doesn’t explain amplitude modulation of pupil size with task difficulty*). It has been suggested that the saccade-locked pupil response occurs because of a transient change in luminance [134, 50]. Laurence Stark ruled out this explanation in his 1965 and 1966 papers [48, 49] by varying luminance (i.e., showing a flash of darkness) at variable delays with respect to the time of a voluntary eye movement, and observed that the saccade-locked pupil response effectively quashed the luminance response when they were evoked simultaneously. Furthermore, if the saccade-locked responses were luminance-evoked, the shape and amplitude of saccade-locked responses should be the same during hard and easy runs of our task, because the overall luminance was the same in each. However, the saccade-locked pupil response had a significantly higher amplitude during hard than easy runs. This couldn’t be explained by differences in saccade amplitude, which may affect pupil size by modulating retinal illumination (see *Supplementary Information*. In addition, the shape of saccade-locked pupil response changed dramatically between the 2 and 4 s tasks (Fig. S3, Fig. S6C). This task-timing-dependent deformation was predicted quantitatively by our model (Fig. S6A,B,D), but is not predicted by existing models of the light reflex. Others have suggested that the saccade-locked pupil response is caused by some visual change other than luminance [135, 136], but again this doesn’t explain why the amplitude of the saccade-locked pupil response scaled with task difficulty, despite the stimulus being the same. Stark suggested that the saccade-locked pupil response was due to changes in accommodation arising from switching fixation locations. The eye movements in his study were 20° in magnitude. This explanation doesn’t make sense for the small fixational saccades and microsaccades that were analyzed in our study (see *Methods, Saccade detection and rate estimation*), when depth was virtually identical for different fixation locations. Furthermore, micro-fluctuations in accommodation during fixation don’t predict fluctuations in pupil size [137, 138, 105]. The function of the saccade-locked pupil response, if there is one, remains unclear [18], however, its origin is more clear. It does not seem to be visually or accommodatively evoked (although see ref [134]). Its causal link to activity in SCi [46, 26, 45] is the strongest evidence we currently have about its control.

### 3.3 Conclusions

It has long been known that the size of the pupil reflects cognition and arousal [1, 13, 40, 18, 21]. However, it was discovered only recently that pupil size and saccades share a common neural input [46, 26, 45], prompting a revision of existing models of the task-evoked pupil response. In the model we propose here, a common, noisy input (i.e., generator function) drives both pupil size and saccades and a gain subsequently modulates pupil size. We find that the common input reflects a cognitive representation of task structure (including its timing), and that the gain reflects arousal level. We offer an algorithm, based on our model, that estimates both components from measurements of pupil size and saccades. Our code toolbox is available at [link].

## 4 Methods

### 4.1 Observers

Observers (N = 10, 6 females, 4 males) were healthy human adults with no known major neurological disorders, and with normal or corrected-to-normal vision. Five observers (O1-5) participated in the 4 s task. Five additional observers (O6-10) participated in both the 2 and 4 s task. All observers were naive to the purposes of the experiment except O1 and O6 (two of the co-authors). O2, O3, O4, O7 and O9 had little to no experience participating in visual orientation-discrimination experiments, while the remaining observers had considerable experience. Experiments were conducted with the written consent of each observer. The experimental protocol was approved by the University Committee on Activities Involving Human Subjects at New York University.

### 4.2 Equipment and setting

Stimuli were displayed on a CRT monitor (22” diagonal; Hewlett-Packard p1230) with a refresh rate of 75 Hz and resolution of 1152×870, custom calibrated for gamma-correction. The task and visual stimuli were controlled with Matlab software based on the MGL toolbox (gru.stanford.edu/mgl). The observer’s eyes were 57 cm from the monitor and head was fixed in a chin-and-forehead rest, which minimized head movement. Pupil area was recorded continuously during task performance using an Eyelink 1000 infrared eye tracker (SR Research Ltd., Ontario, Canada) with a sampling rate of 500 Hz. Calibration was performed before each run of 75 trials to ensure proper measurement of eye position. The room was dark and the door remained shut for the length of the experiment, ensuring dark adaptation was not interrupted. Cell phones were taken away to prevent vibrations and light emissions that might alter engagement or arousal.

### 4.3 Task design

Observers performed a two-alternative forced-choice visual orientation-discrimination task, which varied in difficulty level in alternate runs of trials. We instructed observers to report whether a small, peripheral grating was tilted counter-clockwise or clockwise of vertical by pressing the “1” or “2” keys on a keyboard with the non-dominant hand. Auditory feedback provided immediately after button press indicated accuracy (high tone, correct; low tone, incorrect). Observers were asked to fixate at a central cross (width, 0.7°) and to minimize blinking. The (equiluminant) color of the fixation cross was red on hard runs and green on easy runs to explicitly inform the observer of the current task difficulty. We instructed observers to be as accurate and as fast as possible in responding, and to maintain fixation at all times.

The experiment consisted of 2 sessions, conducted on 2 separate days. Each session comprised 5 separate runs (75 trials/run), alternating between easy and hard. Therefore, observers performed either 3 easy runs for the first session and 3 hard runs for the second session, or vice versa. This order was randomly chosen. 5/10 observers did 2 easy runs in their first session. Observers could rest between runs for as long as they needed, but typically rested for less than 1 minute. Each run consisted of 75 4-second trials, lasting for 5 minutes each, for a total of 60 minutes per session (with breaks).

Each trial comprised a stimulus presentation of 0.2 s followed by an inter-stimulus interval of 3.8 s, in the 4 s version of the task, or 1.8 s, in the 2 s version. During the first 1 or 2 s, respectively, of the inter-stimulus interval, the observer could respond with a button press. If observers missed that response window, no tone would play, indicating a missed trial. Observers missed the response window exceedingly rarely (15/6900 trials, i.e., 0.22%].

Before beginning the experiment, observers were trained initially for 75 or 125 trials depending on their familiarity with orientation discrimination tasks. Orientation threshold was measured as the staircase value on the final training trial and was used to set the initial stimulus tilt in the first hard run of trials in the experiment.

### 4.4 Stimulus

The stimulus was a grating (diameter 1.5°, spatial frequency, 4 cycles/°; contrast, 100%) multiplied by a circular envelope (diameter, 1°) with raised-cosine edges (width, 0.25°). The stimulus had a mean luminance (over space) that was equal to the background luminance (mid-grey), so that it would not be expected to evoke a luminance response from the pupil, driven by intrinsically-photosensitive retinal ganglion cells with large spatial integration areas (i.e., wide dendritic trees [139]). The stimulus was presented in the lower right hemifield, 5° from the center of the screen (Fig. 1C). On easy trials, the tilt of grating was fixed at +-20° from vertical, yielding 93% correct discrimination accuracy on average across observers. On hard trials, the tilt of the grating was controlled adaptively according to prior performance. Specifically, the absolute value of the tilt was controlled by two interleaved 2-down-1-up staircases with initial thresholds of 5° and 0°, respectively, and an initial step size of 1, which converged to 70% discrimination accuracy (on average). The tilt’s sign (i.e., clockwise or counter-clockwise of vertical) on each trial was determined randomly. The staircase value (stimulus tilt) on the final trial of a run was carried over to the following run or session.

### 4.5 Pupil data pre-processing

Pupil size was recorded as the area of a model ellipse in the arbitrary units specified by the Eyelink eye tracker’s firmware. The Eyelink units are proportional to pupil area (mm^2^) (Hayes, & Petrov, 2015). Therefore, it can be easily used in regression models, and that if the pupil size (AU) in one condition is doubled compared to another condition, that relation holds in physical units. Gaze position and pupil area time series were linearly interpolated during and 150 ms before and after blinks, following refs [81].

For estimating the saccade-locked pupil response, we first band-pass filtered the pupil area time series to remove low frequencies that would confound deconvolution, as well as high frequencies that were not physiologically plausible (i.e., measurement noise). For all other analyses (e.g., for data analysis and model fitting), we just low-pass filtered the pupil size signal to remove high-frequency measurement noise. The band-pass filter was a Butterworth 4th order zero-phase filter (“filtfilt” in Matlab) with 0.03 Hz and 10 Hz cut offs. The lowpass filter was a Butterworth 2nd order zero-phase filter with a 10 Hz cut off.

We employed a custom convolution boundary handling method to ensure that we did not generate large, artifactual signals at the edges of the filtered time series. Boundary handling was performed differently for the beginning and end of the time series. The first sample of the time series was repeated N times and concatenated to the front of the time series. The last N samples were mirrored and concatenated to the end of the time series. N refers to the length of signal. This method was employed because the dynamics of pupil size *during* the task (e.g., padding with a mirror image of the signal) are not a good estimate of what was happening *before* the task and in the first few seconds of the first trial. Conversely, the dynamics at the end of the task are likely a better model for what will continue to happen during the next hundreds of milliseconds after the task ends.

We downsampled the pupil time series just for deconvolutions to speed up the compute time. Every 8th sample from the time series was preserved, respecting the Nyquist frequency (this was after the data were low-pass filtered with a cut-off of 10 Hz).

### 4.6 Saccade detection and rate estimation

Saccades (including microsaccades) were detected with the method of Engbert and Mergenthaler [140]. A velocity cutoff of 8 °/s and a duration cutoff of 7 ms were used as inclusion criteria for saccades. The distribution of saccade amplitudes is unimodal, which has led to varying operational definitions of a microsaccade [103]. Our cutoff includes ballistic eye movements commonly called saccades, as well as microsaccades. We refer to both of these as “saccades” throughout. We included both microsaccades and saccades, which are known to have different properties (e.g., the former is involuntary but the latter is voluntary) [141, 142], because previous studies show that a similar constrictory pupil response is observed following either [48, 49, 50, 52, 51].

We quantified the impact of including larger saccades (i.e., greater than 1.5° in amplitude [142]) on our results by repeating our model fitting for each observer and excluding any saccades above 1.5° in amplitude from all steps. The stimulus was positioned 5° diagonally from fixation and the screen edge was 20° horizontally and 15° vertically from the screen edge. Therefore, larger saccades might evoke a pupillary light reflex. The proportion of saccades larger than 1.5° was 3.12% (average across 15 data sets), i.e., quite rare. The median increase in *R*^2^ across observers before vs. after including an upper threshold on saccade amplitude of 1.5° was 0.3% (MAD, 0.7%; N = 15 data sets). The median increase in gain was 0.00011 (mad, 0.00034; N = 15 data sets)). Thus, the impact of larger saccades on our model fitting was negligible, and we obtained comparable results with or without them for our chosen task.

### 4.7 Estimation of the saccade-locked pupil response

For each run of trials, we estimated the saccade-locked pupil response using deconvolution, a common method for determining the finite impulse response function (IRF) of a linear system. We used an algorithm similar to the one used by Knapen et al. (2016) [50], in which we defined a deconvolution design matrix (i.e., a Toeplitz matrix). The first column contained ones at the onset times of each saccade and zeros everywhere else. Successive columns were shifted by one sample. We estimated the impulse response function by inverting the linear system, given the entire measured pupil size time course. We assumed a 4 s duration for the saccade–locked pupil response, which was sufficient to capture the whole response according to informal comparisons of different durations and previous studies [50].

### 4.8 Linear-nonlinear linking model

We used a linear-nonlinear (LN) linking model to predict pupil size from saccade rate and to estimate arousal level. The system’s input, the “generator function,” was assumed to be a time-varying signal corrupted by additive Gaussian noise with standard deviation *σ* equal to 1. The system has two “channels” (Fig. 1B), one leading from the generator function to saccades (Fig. 2A) and the other leading from the generator function to pupil size (Fig. 2E).

We can express the generator function on a single trial as

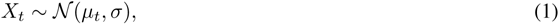

where *X_t_* is the generator function, *x* represents the possible values it can take, *t* is time from trial onset to end, d*t* is determined by the sampling rate of the eye tracker (i.e., d*t* = 1/500 s for our data), and *µ_t_* is the expected value of the generator function (i.e., across trials).

The channel leading from the generator function to saccades contains two operations, the first a threshold non-linearity and the second a post-saccade refractory period. The threshold non-linearity simply subjects the noisy generator function to a fixed threshold, represented in Fig. 1B as a step function. Any time the generator function rises above the threshold is represented by a one or otherwise by a zero in a binary sequence. This binary sequence represents the time course of saccades (and ignores saccade amplitude). The threshold is implemented by mapping trial-averaged saccade rate through a non-linear function (the inverse cumulative distribution function of a Gaussian with standard deviation *σ* = 1). The threshold non-linearity is similar to an inhomogeneous Poisson process (see *Supplementary Information, Equivalence of our model with an inhomogeneous Poisson process*). The second computation, the post-saccadic refractory period, models the phenomenon of inhibition in saccade generation after a saccade has just occurred as a multiplicative scaling of the entire trial-averaged saccade rate function (see *Supplementary Information, Modelling the post-saccade refractory period*). We adopted a statistical model called a renewal process to account for the refractory period, by starting with a Poisson process and scaling the rate by 1/*k* — equivalent to preserving every *k*-th saccade [143]. The Poisson process has an exponential distribution of inter-saccade intervals but the renewal process has a gamma distribution. This channel of the model yields an (adjusted) trial-averaged saccade rate function. We can therefore write the mapping between saccade rate and the generator function as:

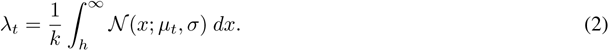

*λ_t_* is saccade rate (a probability between 0 and 1), *k* is the shape parameter of a gamma distribution fit to the inter-saccade intervals, and *h* is a fixed threshold (arbitrary, but fixed to 1 in our model fitting and simulations).

The channel leading from the generator function to pupil size contains two operations, a multiplicative gain (i.e., an amplifier) and a low-pass filter (i.e., a temporal integrator). The gain simply scales the noisy generator function by a number *g*. We assume that gain is constant within a trial and may change across trials, but that its expected value across trials is *g*. The model’s prediction of the task-evoked pupil response is equal to the circular convolution of the linear filter (see *Methods, Estimating the linear filter*) with the expected value of the mean-subtracted, gain-modulated generator function, plus an additive offset. We used circular convolution to capture the periodicity of the generator function in our task, in which the inter-trial interval was fixed. Mean-subtracting the generator function before the gain corresponds with a particular sort of gain modulation, in which the DC offset of the generator function doesn’t change with gain (like a stereo amp might do); only its amplitude (distance from peak to trough) changes. The model fits data equally well without mean-subtracting the generator function, but not mean-subtracting makes three unjustifiable assumptions: (1) that the gain and DC offset of pupil size are strongly (positively) correlated, (2) that the DC offset is (much) higher than the mean pupil size, and (3) that the threshold depends on gain. Instead, we simply mean subtract the generator function and later add to the model prediction an additive offset *b*, equal to the mean pupil size over a run of trials. This second channel of the model yields a prediction of trial-averaged pupil area (i.e., the task-evoked pupil response). We can write the relation between the task-evoked pupil response and the expected value of the generator function as

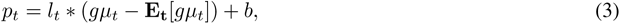

where *l_t_* is the linear filter, *g* is the gain, *b* is the additive offset, “*” denotes circular convolution, and **E** denotes an expectation. The mean and standard deviation of the gain-modulated generator function are:

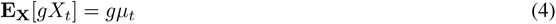

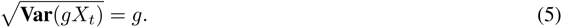

Therefore, both the amplitude and standard deviation of the generator function scale with gain.

### 4.9 Parameter estimation

We estimated the generator function and gain using trial-averaged pupil and saccade data. We colloquially refer to this estimation method as “ascending one channel of the model and descending the other,” because we started with measured saccade rate, estimated the expected value of the generator function, and finally predicted the trial-averaged pupil response, which can be compared to the measured pupil response (Fig. 2). This procedure yielded estimates of the expected value of the generator function and three parameters — gain, offset, and *k* — of which only gain was fit separately for each run.

To estimate gain, we performed a linear regression between the measured and predicted trial-averaged pupil response. Gain was equal to the beta value of this regression; i.e., it simply scaled the prediction. The predicted pupil response was computed as the circular convolution of the linear filter and the mean-subtracted and amplified expected value of the generator function, plus an additive offset (mean pupil size over a run) (Eq. 3).

To compute the expected value of the generator function, we mapped the adjusted saccade rate function (i.e., “adjusted”: times *k*, see below) through the inverse function of a cumulative Normal distribution. Saccade rate at time *t* was assumed to be the integral of a Gaussian with standard deviation *σ* = 1 beyond a fixed threshold (i.e., from the threshold to infinity). The mean of this Gaussian was the expected value of the generator function at time t. The threshold was always assumed to be 1, but its value is arbitrary as long as it is fixed. We can write the expected value of the generator function *µ_t_* as a function of saccade rate *λ_t_* by inverting Eq. 2:

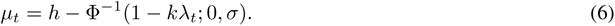

Φ is the cumulative probability function for a Gaussian distribution and Φ*^−^*^1^ is its inverse. We use Eq. 6 to estimate the expected generator function *µ_t_*.

The entire transformation between trial-averaged saccade rate and pupil area can be expressed as

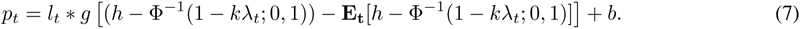

Note that there is a non-linearity in between saccade rate and pupil area: the inverse cumulative Gaussian.

We estimated *k*, the parameter controlling the post-saccadic refractory period, for each observer by fitting a gamma distribution to the empirical distribution of inter-saccadic intervals (for justification, see *Supplementary Methods, Modelling the post-saccade refractory period)*. *k* is the shape parameter of a gamma distribution. For each run, we multiplied the saccade rate function by *k* to compensate for the reduction in saccades caused by the refractory period, equivalent to preserving only every k-th saccade [143]. The multiplier *k* on saccade rate scales the input to the non-linearity Φ, meaning that *k* influences pupil size in a non-linear way (Eq. 7). In practice, however, including *k* in the model had a negligible effect on the model’s parameter estimates and goodness of fit (see *Supplementary Information*). Thus, although modelling the refractory period was theoretically justified, it was practically unnecessary for real data. This was true under the assumption of a gamma renewal process or under the assumption of an absolute post-saccadic refractory period of 125 ms [144].

We confirmed that our model fitting procedure was able to recover the gain and generator function, when specified *a priori*, with parameter recovery simulations (see *Supplementary Information, Parameter recovery: our model*).

### 4.10 Estimating the linear filter

The linear filter was estimated by fitting a parametric form to the deconvolved saccade-locked pupil response (Fig. S3A). We optimized for the best combination of parameters (*n* and *t*_max_) that best described the saccade-locked pupil response as a gamma-Erlang function, a common approach in the pupil modelling literature [43, 50, 42]. We used the fminsearch function (Nelder-Mead) in Matlab for the optimization. The equation for the parametric form was:

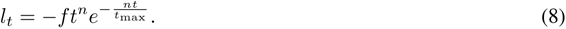

The initial points for *n* and *t*_max_ were randomly drawn from the intervals (0,20) and (600,4000), respectively. These initial points were chosen based on the parameter range used in previous studies [43, 50, 42]. Maximum function evaluation and maximum iteration both were set to 5e4. One set of parameters were fitted for each observer. We set *f* (the scale factor) such that the parametric form had the same height (minimum) as the measured saccade-locked pupil response, equivalent to multiplying the estimated filter by a normalization factor (height of filter estimate divided by height of saccade-locked pupil response). This meant that estimates of gain were relative to the amplitude of each observer’s saccade-evoked pupil response. Therefore, gain estimates were in the same range for all observers and gain modulation (i.e., with task difficulty) could be compared directly across observers (Fig. 3C). This normalization was a crucial pre-processing step [145] because foreshortening error varies for each observer due to the angle of the eye tracker (i.e., adjusted for each observer’s height), which affects the range of measured pupil response amplitudes (and thus, gains, also). Note that this is different from divisive baseline correction [86, 42, 146], in which the pupil size signal is divided by its mean in order to account for a supposedly nonlinear interaction between the signal’s amplitude and additive offset. We did not perform divisive baseline correction on our data because the task-evoked pupil response has been shown to scale linearly with baseline within normal range (i.e., not at the highest or lowest pupil sizes, where there must be compressive nonlinearities due to biomechanical limits) [86]. We set *f* based on the minimum of the saccade-locked pupil response because the alternative, regression, caused mis-estimation of the filter’s amplitude and other parameters due to distortion of the saccade-locked response (caused by the threshold non-linearity). Setting *f* in the way that we did avoided this, according to our simulations (see below).

The time-to-peak (*t*_max_) and width of this filter (*n*) varied per observer, and these idiosyncrasies were critical for explaining variability in the dynamics of the task-evoked pupil response for some observers (see *Supplementary Information, Individual differences in saccade rate and linear filter were sometimes critical to accurately predict pupil size*), as Parker & Denison (2020) also suggested [42].

We used an (upside-down) gamma-Erlang function as a parametric form for three reasons. First, our assumption was that the linear filter is low-pass and constrictory. Second, a gamma-Erlang function has been shown to model the aggregate input-output relationship of a system with cascades of exponential computations with different time constants (e.g., a multi-synaptic neural circuit like the parasympathetic pathway of the pupil) [43]. Third, parameter recovery simulations revealed that this parametric form eliminates a large amount of bias in the filter estimate relative to a ground truth filter (see *Supplementary Information, Justification for linear filter estimation method*).

### 4.11 Model comparison

To assess model generalizability we computed the reduction in *R*^2^ between the in-vs out-of-sample fits as well as ASI and HSI indexes (see *Results* for details). For our model, we computed the average gain across all easy and hard runs separately for each observer, and used these average parameter values for the out-of-sample model predictions. For the consensus model, we did the same, but with the best-fit amplitudes for the impulses at trial onset, button press, and for the time-on-task boxcar for the easy and hard data. The duration of the time-on-task boxcar was from trial onset to the mean reaction time. To fit the consensus model, we used the PRET toolbox [42] to implement a model similar to that proposed by Denison and colleagues (ref [42]) and others [34, 50, 36]. These models are variants of one another, but share the same overarching assumptions: a linear model with impulses near trial events as input and a dilatory linear filter. We didn’t fit the latency of the impulses from the task events, as Denison and colleagues suggest doing, because it was not standard and would afford the model too much flexbility (also see *Supplementary Information, Parameter recovery: deconvolution method of Wierda and colleague*). When we did fit these latencies, the in-sample fit of the model improved and the generalizability decreased, consistent with the idea that it made the overfitting worse.

For the consensus model only, the computed ASI and HSI indexes were sometimes extremely large or small because one of the best-fit amplitudes would go to nearly zero on one data set and would be larger than zero for the other dataset. This suggested that the three parameters traded off in the consensus model, again suggesting it overfits.

## 5 Acknowledgements

We thank Mike Landy, Rachel Denison, Eero Simoncelli, and Nick Steinmetz for their comments on the research and Elisha Merriam and Nastaran Arfaei for their help with early versions of the experiments. This research was supported a National Eye Institute grant (R01-EY025330) to D.J.H. and a National Defense Science and Engineering Graduate fellowship to C.S.B.

## 6 Author Contributions

Conceptualization, C.S.B., S.M., and D.J.H.; Methodology, C.S.B., S.M., and D.J.H.; Software, C.S.B. and S.M.; Formal Analysis, C.S.B. and S.M., Investigation, C.S.B. and S.M.; Writing - Original Draft, C.S.B.; Writing – Review & Editing, C.S.B., S.M., and D.J.H.; Visualization, C.S.B.; Funding Acquisition, D.J.H; Supervision, D.J.H.

## 8 Supplementary Information

### 8.1 Supplementary Appendix

### 8.2 Goodness of fit depended on gain

Model fits were significantly better on runs when gain was higher (Fig. 3F), suggesting that our model fit better when arousal level was higher. The relationship between gain and *R*^2^ was strongly heteroscedastic. When gain was extremely low (i.e., near zero), *R*^2^ was also very low, near zero. For a middle range of gain values (but below the mean gain value), *R*^2^ was highly variable, ranging from near 0 to near 1. When gain was high (above the mean), *R*^2^ was highly concentrated just below 1, and only deviated below 0.5 once (Fig. 3F). We ran a regression model, fitting *R*^2^ with gain, including difficulty and the interaction between gain and difficulty as covariates, and accounting for systematic variations in gain across sessions and observers (see *Supplementary Information, Statistical analysis*). Gain remained a significant predictor of *R*^2^, meaning that *R*^2^ was higher when gain was higher overall across observers despite the inclusion of these other factors (F(139) = 29.82, p = 2.12e-07, N runs = 143). This suggests that at low arousal, influences on pupil size other than those specified by our model may dominate. If there is any late noise in the channel connecting the generator function to the pupil (i.e., noise in iris muscles or ciliary ganglion), as gain decreases, the ratio between generator noise and late noise will drop, saccades and pupil size will become increasingly de-correlated, and the *R*^2^ of the model fits will go down. Hence, this result is evidence that there is late noise whose influence on pupil size is usually small, but can dominate when arousal is very low. It could also be due to variable sampling error in the estimate of the generator function (i.e., due to the number of saccades, which varies from run to run), however our parameter recovery results speak against this possibility.

### 8.3 Individual differences in saccade rate and linear filter were sometimes critical to accurately predict pupil size

Individual differences in saccade rate (Fig. S2) and linear filter (Fig. S3A) predicted idiosyncrasies in the shape of each observer’s task-evoked pupil response (Fig. S4). We quantified the contribution of individual differences in saccade rate to the goodness of fit of the model by using a grand-mean saccade rate function across all observers to re-fit the model to each observer’s data. Note that the filter was still individualized. We computed separate grand mean rate functions for easy and hard runs. The mean reduction in *R*^2^ between the full model (i.e., with individualized saccade rate functions for each run) and the reduced model was 14.51% (sem, 4.90%; N = 87 runs). Some observers had large reductions (e.g., O3, 55.19%, O6, 43.58%), suggesting that individual differences in the rate function can be crucial for explaining idiosyncracies in pupil size. We performed a similar model comparison, but used a grand mean linear filter (across all observers) for each observer’s fit (and saccade rate was individualized). The mean reduction in *R*^2^ was 11.34% (SEM, 4.35%, N = 86 runs). For observer O9, there was a substantial reduction in fit (39.11%), demonstrating that it can sometimes be critical to individualize the linear filter. For observer O3, there was an unexpected enhancement in fit (−22.81%). Notably, O3’s filter estimate was very delayed and broad compared to the others (Fig. S3A, O3), suggesting that it may have been an inaccurate filter estimate due to issues in data collection (O3 wore eye glasses which could have disturbed the eye tracking signal during eye movements). On average, individual differences in the linear filter and saccade rate accounted for only 10-15% of the variance in the task-evoked pupil response. That said, we still recommend individualizing because it can be unexpectedly important for specific observers.

### 8.4 Saccade amplitude doesn’t explain amplitude modulation of pupil size with task difficulty

The pupil responds most strongly to light. Hence it is natural to ask whether our pupil response measurements were confounded by the pupillary light reflex [10]. There was a large change in luminance at the screen’s edge in our experimental set-up. Hence, difficulty-dependent differences in the frequency and/or amplitude of eye movements might have caused a difference in retinal illumination and therefore a difference in pupil responses. Saccade rate was largely indistinguishable for easy vs. hard trials, so we focused on saccade amplitude.

If observers simply made larger saccades during hard trials, that may explain difficulty-dependent modulation of pupil size. We found that for a majority of data sets, there was no significant difference in saccade amplitude between easy and hard trials (t-test: p > 0.05), despite clear amplitude modulation of the saccade-locked (Fig. S3B, Fig. S6C) and task-evoked pupil responses (Fig. S4) with task difficulty. For 7/15 data sets, we found significantly higher saccade amplitude for hard than easy trials (t-test: p < .05). These differences were, however, extremely small: Cohen’s d ranged across observers from 0.05 to 0.11, and barely evident in histograms of saccade amplitudes (i.e., for all 15 data sets, the distributions for hard and easy were strongly overlapping). For this statistical test, we log-transformed the saccade amplitudes, as they were heavily right-skewed, and this made the data roughly Gaussian. In summary, larger saccade amplitude could not account for larger pupil responses for hard than easy trials, suggesting that the saccade-locked pupil response is not a luminance response (see *Discussion* for additional arguments for why a luminance artifact cannot explain saccade-locked and task-evoked pupil responses).

### 8.5 Supplementary Methods

### 8.6 Justification for linear filter estimation method

The true linear filter, which captures the relation between the generator function and pupil size, cannot be directly estimated because we do not directly measure the generator function. Because the mapping between saccades and pupil size is non-linear (Eq. 7), the saccade-locked pupil response is a deformed (i.e., biased) version of the true filter, under our model assumptions. We performed parameter recovery simulations to characterize this deformation and to determine its practical impact on our model fitting. Assuming a ground truth filter (a gamma-Erlang function) and generator function (estimated from saccade data from one of our observers) and some value for gain, we simulated 75 4-second trials of pupil size and saccade data, and then deconvolved the saccade-locked pupil response. In our simulation, i.i.d. normally-distributed noise was added to the generator function on each trial and saccades were simulated as the times of threshold crossings of the noise-injected generator function. We repeated the simulation a number of times, changing the parameters of the gamma-Erlang filter, changing gain, and swapping in generator functions estimated from different observers’ data. The deformation of the (simulated) saccade-locked pupil response from the ground truth filter was highly consistent and had two notable features. First, the recovered IRF was biphasic, exhibiting a late dilatory component not present in the ground truth (Fig. S6A). Second, the width around the minimum was larger in the recovered IRF. In spite of this, the time-to-minimum was preserved. Furthermore, changing the ground truth gain led to precise multiplicative scaling of the saccade-locked response. For example, doubling gain led to a doubling of the minimum of the saccade-locked response (Fig. 3H, Fig. S6B-D, Fig. S3B). That suggests that the deformation doesn’t affect scaling of the response with gain, nor its time-to-minimum. We decided to fit a monophasic parametric form (gamma-Erlang) that would eliminate the first aspect of the deformation, the late dilatory component. For the second aspect of the deformation, the increase in response width, we refit our model on the real data using an artificially lowered response width for each observer’s filter. We lowered the width by an amount that made the estimate and ground truth match in the parameter recovery. We found that the resulting influence on the goodness of the fit of the model was negligible. In summary, although the saccade-locked pupil response is technically a biased estimate of the linear filter due to the threshold non-linearity in between saccades and pupil size, our simulations revealed that this bias is not large and that our parametric fit eliminates a substantial portion of it (Fig. S6A, Fig. S3A). Nonetheless, we see value in developing methods to estimate the linear filter from saccade and pupil size measurements that takes into account the deformation.

### 8.7 Parameter recovery: our model

We assessed model identifiability by generating synthetic data from our model (i.e., assuming a ground truth gain and generator function) and applying our algorithm to estimate the ground truth model parameters (Fig. S12). Accurate recovery of the ground truth parameters would indicate an identifiable model. We were also interested in determining the amount of data needed to accurately estimate the parameters and achieve good fits, so we additionally varied the number of simulated trials. We generated the ground truth generator function from the grand mean saccade rate function across all hard correct trials (N = 3450 trials, N = 10 observers, 4 s task; Fig. S12A). We did this because we wanted the ground truth we chose to reflect the true expected value of the generator function as much as possible (i.e., to minimize sampling error). The ground truth filter was a gamma-Erlang function with parameters close to those observed in the data (tMin = 900, n = 10, f = -1; Fig. S12B). We assumed a threshold of 1 and *σ* of 1. First, we simulated 75 4-second trials of pupil size and saccade data, matching the amount of data we used to fit the model to the real data on each run (Fig. S12C). We set the ground truth gain to 1 or 2 in different simulations, and used our model fitting procedure to estimate the gain and generator function from the simulated data. The filter estimate and *k* were kept fixed across simulations varying in gain, matching how we analyzed multiple runs of real data.

The ratio between the recovered gain values (before and after doubling the ground truth) was 1.998 (Fig. S12D), suggesting that even with 75 trials of eye data, our algorithm can accurately recover modulation of gain. The *R*^2^ of the fit to the synthetic data was 95.85%. Both findings were impressive given that the synthetic saccade rate function (i.e., from the simulated threshold crossings, N trials = 75) was very undersampled relative to the original (using 3450 trials) and had numerous time points where the rate reached zero. Correspondingly, the recovered generator function was noisy and a poor estimate of the ground truth (average d’ over time between ground truth and estimate, 0.57; max d’, 2.64; Fig. S12A). This suggests that more than 75 trials is necessary to accurately estimate the generator function. Taken together, these results demonstrate that an accurate estimate of the expected generator function is not necessary to accurately estimate gain or predict pupil size.

Next, we repeated the procedure but increased the number of simulated trials to 3450 to match the amount of input data. The estimated modulation of gain was still very accurate (ratio of estimated gains, 2.05; *R*^2^, 0.981; Fig. S12C,D), and additionally the estimated generator function was now very accurate as well, with an average d’ (over time) with the ground truth of 0.04. The generator function was recovered accurately except for time points where the synthetic saccade rate was undersampled due to chance, i.e., when saccade rate was zero (max d’ during these periods, 1.09; Fig. S12A). This led to brief troughs in the recovered generator function that varied in amplitude across repetitions of the parameter recovery simulation (i.e., across random draws from generator noise distribution), meaning that they were unreliable. This suggests that thousands of trials are necessary to accurately estimate the generator function’s dynamics. That said, sharp troughs in the generator function like these were not observed in the real data when using similar amounts of trials (Fig. 4A,C-E, Fig. S5B, Fig. S8, Fig. S9A). This suggests that there may be some attribute of the saccade timing we didn’t model in the data that protects against these variable troughs, i.e., 3450 trials may suffice for accurate estimation. The recovered generator function was invariant to the ground truth gain, suggesting that gain and the generator function were distinct and identifiable. For both amounts of synthetic data (75 and 3450 trials), the estimated filter was deformed as expected (see *Methods, Estimating the linear filter*; Fig. S6; Fig. S12B).

### 8.8 Parameter recovery: deconvolution method of Wierda and colleagues

We performed parameter recovery for the deconvolution method of Wierda and colleagues [44], which is based on a similar method proposed by Hoeks and Levelt [43]. The model of Wierda and colleagues assumes (1) that the input to the pupil is a sequence of variable-amplitude “attentional pulses” that occur every 100 ms, with no input at any other moment in time, and (2) that pupil size is the convolution of this input with the dilatory filter specified by Hoeks and Levelt. Wierda and colleagues propose a deconvolution method for estimating the pulses from trial-averaged pupil responses, given the assumption that the pulse amplitudes must be positive. Our parameter recovery procedure consisted of setting a ground truth input (“attentional pulses”), simulating the trial-averaged pupil response by convolving this input with a filter [43], and then estimating the input using deconvolution. This procedure allows us to test whether the proposed deconvolution method can accurately recover the input parameters, when they are known. If not, then the method is underconstrained and therefore not suitable for use with real pupil data. We used the code provided by Wierda and colleagues in their paper, and adopted all the same parameters and procedures, so we refer the reader to their methods and code for details [44]. We generated two different types of ground truths - one was Gaussian white noise (mean, 0.4; sd, 0.1) with a similar mean, range, and standard deviation as the best-fit input from their experimental data, and the second was a series of two impulses like the input form assumed in Hoeks and Levelt [43]. The Gaussian input was a sparse vector (spacing between non-zero elements, 100 ms), where every impulse with non-zero amplitude was drawn from a Gaussian distribution. We quantified the goodness of fit of the assumed vs. predicted pupil response with *R*^2^ and also measured the recovery of the ground truth input with *R*^2^. For the Gaussian noise input, across 10 different draws of noise, the average *R*^2^ of the pupil response prediction was 99.99% (SEM, 6.20e-04), but the average *R*^2^ describing the recovery of the input was 24.42% (SEM, 4.18). For the Hoeks-and-Levelt-like input composed of two impulses at 1 and 1.5 s into the trial with the same amplitude, the *R*^2^ between pupil ground truth and prediction was 99.87%, and the *R*^2^ between the ground truth and recovered input was 20.45%. The parameter recovery failed, despite the model’s perfect fits to pupil data, demonstrating that the deconvolution method of Wierda and colleagues is underconstrained.

### 8.9 Pupil noise analysis and simulation

We measured trial-to-trial variability in pupil size (“pupil noise”) during task performance and analyzed its dynamics and modulation with task difficulty (Fig. S7). We performed the same analyses on simulated pupil data to validate our approach. For each observer, we aligned the pupil responses from each trial by subtracting a constant so that each response started from zero (i.e.,“epoching”). We computed the pupil noise level as the standard deviation of pupil size across trials at each point in time from 0 to 4 s. Epoching pupil data in this way arbitrarily sets the standard deviation across trials to zero at trial onset and forces it to remain low at first, hence the characteristic increasing shape of our functions. We were interested in large deviations from this increasing shape, which would suggest non-stationary noise over time (Fig. S7A). And we were interested in multiplicative scaling of this time-varying function with task difficulty, which would suggest modulation of a gain (on the noise level) that is static within a trial (i.e., our model assumption).

To validate our analysis approach, we simulated 375 trials of pupil data from our model, using a simulated generator function and filter originally estimated from O3’s data, and performed the same analyses on the simulated data. We picked O3, because their saccade rate was similar for easy and hard runs, but gain was 1.84x larger (N = 375 trials, best-fit gain, 0.0144 for hard runs, 0.0078 for easy). Noise modulation, measured as the ratio of the average pupil noise level over time for hard vs. easy, was 1.19 (data) and 1.84 (Fig. S7A, “simulation”). Although there was a discrepancy in magnitude (1.19 vs. 1.84), the modulation was in the correct direction and well described by a single multiplier (Fig. 7A). This shows that the ground truth input noise can be stationary within a trial and the estimated standard deviation (over time) will take on this characteristic increasing appearance, analogous to Brownian motion (Fig. S7A, “simulation”). We didn’t observe large deviations from this shape in the data across observers, suggesting the noise is stationary over time.

### 8.10 Modelling the post-saccade refractory period

In our model, it was important to include a post-saccade refractory period because real saccades exhibit an approximately 125 ms refractory period [144], whereas the Gaussian noise added to the generator function could generate saccades in rapid succession with unrealistically short inter-saccadic intervals. Saccades exhibit a refractory period that resembles the one commonly observed in neural spiking, suggesting it is neural in origin [141, 142, 144]. The inter-saccadic interval distribution for microsaccades and small fixational saccades differ [141], with the former following an exponential distribution as expected for a Poisson process (i.e., random timing, fixed rate) and the latter approximating a gamma distribution. Our data, which included both microsaccade and small saccades, should therefore be somewhere in-between. Following Victor and others [143, 147, 148], we modelled the saccadic refractory period as a gamma process, a type of renewal process (i.e., a Poisson process with a refractory period) in which the distribution of inter-event intervals is a gamma distribution with shape parameter *k* [143]. This approach has the special property that it can interpolate between a Poisson process, i.e., with a refractory period of 0 s (*k* = 1) and completely random event times, and a gamma process, i.e., with longer refractory periods and more regular event timing for larger *k*. A common way of simulating a gamma process is to simulate events from a Poisson process and preserve every *k*’th event, i.e., every *k*-th saccade, while deleting the others [143]. The rate of the Poisson process is therefore *kλ* and the rate of the gamma process is *λ*. Therefore, to invert this refractory period (as we would like to do in order to estimate the generator function), we must multiply the measured saccade rate by *k*. In real data, it should be possible to independently estimate *k* by fitting a gamma distribution to the empirical interval distribution.

We performed a simulation to validate this approach and to demonstrate that it generalizes to an inhomogeneous poisson process (i.e., when saccade rate changes over time), when the inter-saccadic interval distribution encompasses all time points from *t* = 0 to trial end (Fig. S13). We started with the average saccade rate function (hard runs only) from observer O1-5 and simulated 50000 trials of saccades from an inhomogeneous poisson process. Next, we preserved every k’th saccade, where *k* was constrained to be an integer: i.e., 1, 2, or 3, within the range that produced inter-saccadic interval distributions that resembled the data (note: *k* = 2 was already too high to resemble most observer’s data, so it was a good upper bound). We computed the intersaccadic interval distribution and saccade rate (i.e., average of saccade counts) with and without the refractory period. The interval distribution was well described by an exponential distribution (*R*^2^ > 0.95) without the refractory period (Fig. S13A) and by a gamma distribution (*R*^2^ > 0.95) with the refractory period (Fig. S13B). The discrepancy between the saccade rate function with and without the refractory period was well described by a single multiplier in a regression (*R*^2^ > 0.98; Fig. S13C,D). For a ground truth *k* of 2, the best-fit beta value was 2.01. The best-fit *k* to the inter-saccadic interval distribution with the refractory period (i.e., the independent estimate of k) was 2.30, suggesting that this estimation procedure can recover the ground truth *k* (*k* must be an integer so the estimate and ground truth will be equal after rounding). Note that preserving every k-th saccade is a nonlinear transformation to the saccade rate, however for the range of saccade rates and *k* values observed in the real data, the nonlinearity was approximated very well by linear scaling.

We performed a second simulation with the same procedure, but we modelled an absolute refractory period of 125 ms, based on measurements of saccade timing [144], by removing any saccades within 125 ms of a prior one. Like the first simulation, this also led to a change in saccade rate that was well described by a multiplier (*R*^2^ = 0.98), despite being a nonlinear transformation. The estimated *k* (2.25) was different than the beta-value (1.21), but this was expected given that the process was not a gamma process. Our model provided a good fit for any multiplier on saccade rate (as long as the saccade probability was *<* 1). This suggests that the model provides a good fit even if the true refractory period does not follow a gamma process, but instead is better modelled by an absolute refractory period. It’s natural to ask whether it is appropriate to use an inhomogeneous Poisson process in our simulations of the impact of the post-saccade refractory period, given that we modelled saccade generation using a threshold nonlinearity instead. We confirmed the equivalence of these approaches in the following section (*Supplementary Information, Equivalence of our model with an inhomogeneous Poisson process*), validating our assumption in the refractory period simulations.

In practice, we found that the influence of including *k* in the model fitting was negligible. The best fit *k* for all observers except O4 and O5 was very close to 1. For O4 and O5, *k* instead rounded to 2 (Table S1). Furthermore, the effect of *k* was locally linear due to being in the far tail of the inverse cumulative Gaussian nonlinearity. Indeed, we found that if we artificially increased or decreased *k*, gain would simply trade off with it (a very small amount), producing an indistinguishable similar goodness of fit (mean difference in *R*^2^, < 0.001) (Fig. S14). This change in gain (0.00025 for O1, 0.00033 for O4) was negligible, more than an order of magnitude smaller (e.g., 25x smaller for O1, 15x for O4) than the difference in average gain between hard and easy runs of trials.

### 8.11 Equivalence of our model with an inhomogeneous Poisson process

It is easy to show that the thresholding in our model is equivalent to an inhomogeneous Poisson process with rate *λ_t_δt*.

At each timepoint, a saccade is either generated or not, with Bernoulli probability

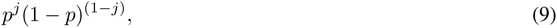

where *p* is the probability of saccade occurrence, equal to *λ_t_δt*, and *j* is 0 or 1 representing whether a saccade occurred or not. *p* is equal to the integral beyond the threshold *h* in the Gaussian distribution of the generator noise (Eq. 2) multiplied by *δt*, the smallest unit of time determined by the sampling rate of the eye tracker.

Over multiple trials, this process is captured by a binomial distribution:

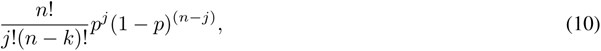

where *n* is the number of events. According to the Poisson limit theorem, as *n* goes to infinity, the binomial converges to a Poisson distribution. We write the probability of a saccade occurring, expanding *p* to *λ_t_δt*:

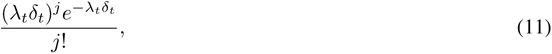

which is the equation of an inhomogeneous Poisson process.

### 8.12 Statistical analysis

We used regression models to fit behavioral performance and task difficulty with our model’s estimates of gain or offset. We used the fitlme and fitglme functions in Matlab to fit linear mixed effects models or generalized linear mixed effects models, respectively. We included random slopes and/or intercepts for each session and/or observer, depending on the analysis, i.e., we allowed the best-fit parameter values to vary by each session and/or observer. This allowed us to account for systematic differences in the range of gain and offset across days or observers, as well differences in their strength of modulation with independent variables.

For fitting *R*^2^ with gain, we ran a linear mixed effects model and included random slopes and intercepts for each observer and session. For fitting reaction time with gain, we did the same but on all the data (easy and hard), and included difficulty level as a covariate. For this model and others, we reported the *R*^2^ of the full model including the fixed and random effects (random slope and intercepts for sessions and observers). For the model predicting reaction time from the three-way interaction of gain, accuracy, and difficulty, we didn’t include random effects because there weren’t enough data to justify it.

For fitting difficulty with gain or offset, we ran a generalized linear mixed effects model (logistic regression), including random slopes and intercepts for each observer and session.

P-values were not provided for the random effects because of the known issue with estimating the degrees of freedom [149]. Instead, we report the p-value of the fixed effects and their interactions (when relevant) with the inclusion of the random effects in the model, and/or just the total *R*^2^ of the model. Some of the data were heteroscedastic (i.e., *R*^2^ and gain) and when we log-transformed the data, making the data more homoscedastic, the null-hypothesis significance tests gave the same result.

For t-tests, we used the ttest2 function in Matlab, which implements a paired t-test.

For distributions of ratios, we report the median as well as the mean, as the ratio of standard normal random variables is known to be skewed and heavy tailed. The median exhibits less bias as an estimator of the ground truth ratio.

---

**Figure 5: Fig. S1.**
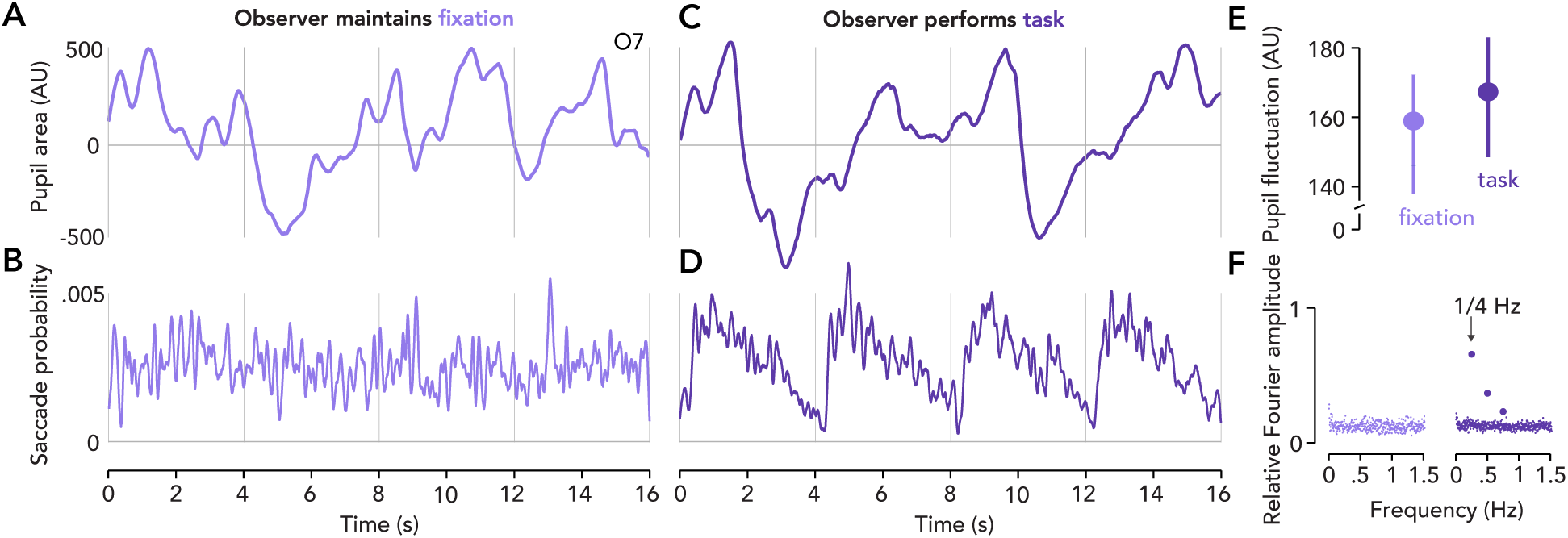
Pupil size and saccade rate during fixation and task performance. We had five observers (O5-10) fixate on a central cross for 10 five-minute runs (on a separate day than the task) to measure pupil size and saccades in the absence of a task. (**A**) Light purple curve, pupil area (mean-subtracted) over 16 s while observer maintains fixation (i.e., no stimulus or task). All data from O7. (**B**) Light purple curve, saccade rate (averaged over 180 16-second sequences) during fixation. Saccade rate is a probability between 0 and 1. (**C**) Dark purple curve, pupil area (mean-subtracted) over four hard trials of the 4 s task. (**D**) Dark purple curve, saccade rate (averaged over 180 sequences of four trials). Fixation and task plots share same ordinate limits. (**E**) Extent of pupil fluctuation during fixation vs. task performance. Pupil fluctuation was measured as the standard deviation of the pupil area signal over a 4 s time bin. Circle, mean pupil fluctuation. Error bar, 95% confidence interval (N datapoints = 720). (**F**) Fourier transform of the entire pupil area time course (frequency response over mean-subtracted run-concatenated data). Light purple, fixation. Dark purple, task. Each dot, amplitude of a particular frequency component. Arrow, emphasizing trial onset frequency of 1/4 Hz. Other elevated dots, harmonics. DC component removed to emphasize other frequencies. Pupil size fluctuates to a similar degree during task performance and rest, however these fluctuations become organized during task performance, entraining to trial onsets. Saccade rate exhibits similar entrainment.

**Figure 6: Fig. S2.**
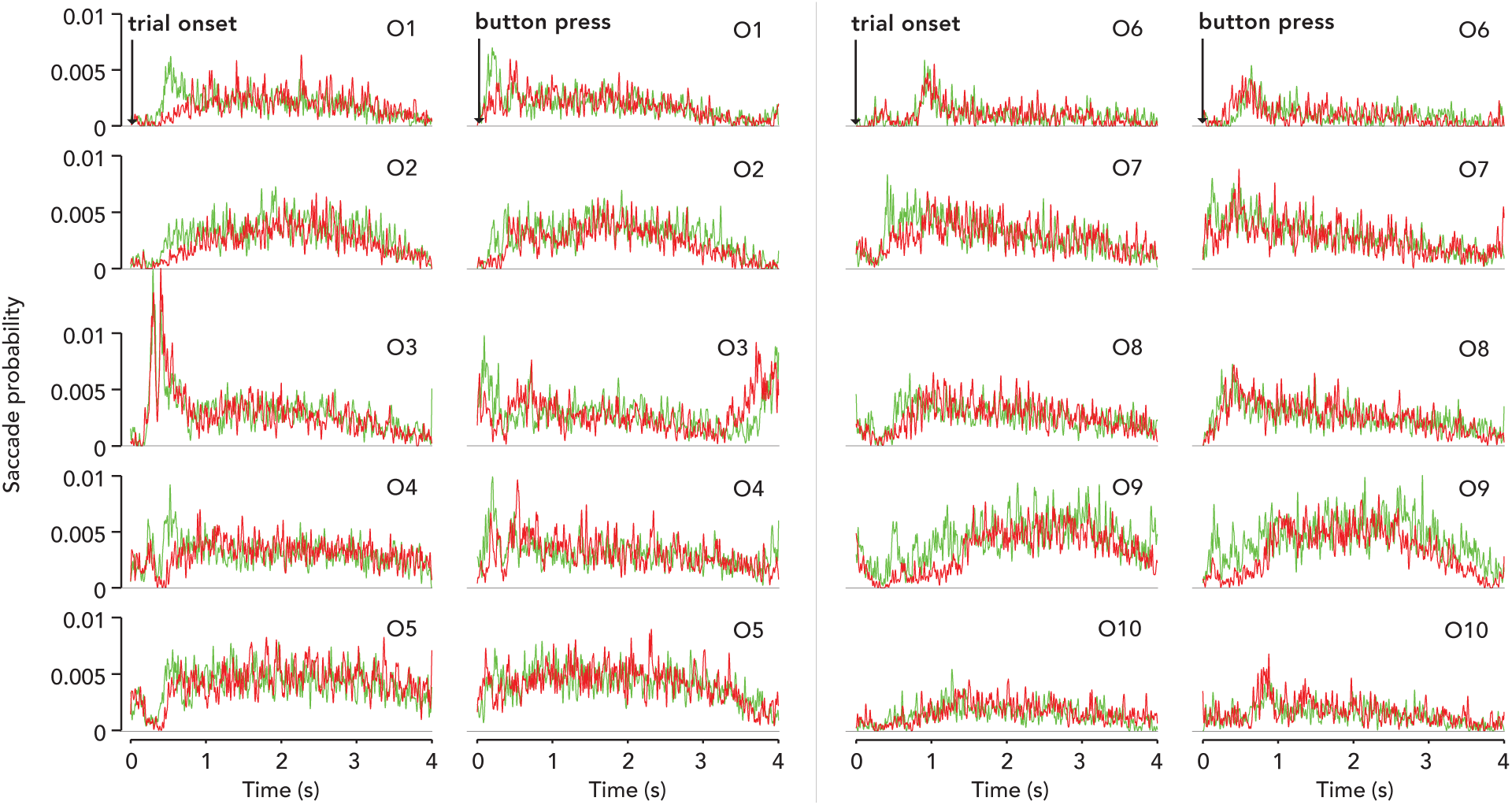
Saccade rate for each observer, time-locked to trial onset or button press. Curves, saccade rate (probability between 0 and 1) over a trial (4 s task). Red, hard runs. Green, easy runs. Left columns, locked to trial onset. Right columns, locked to button press. Each pair of subpanels is a different observer from O1-10.

**Figure 7: Fig. S3.**
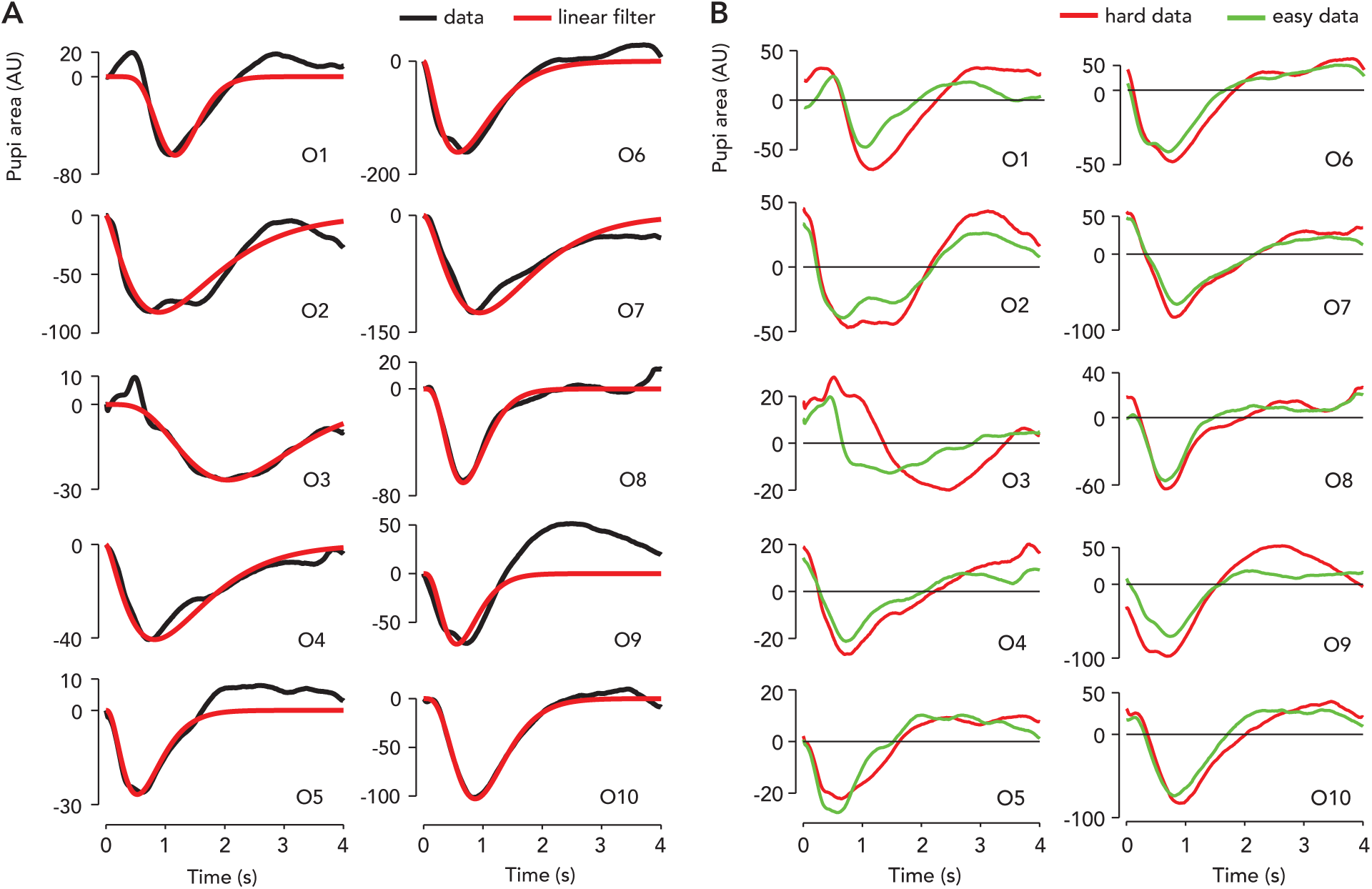
Saccade-locked pupil response and best-fit linear filter for each observer. (**A**) Black curve, saccade-locked pupil response, averaged over ten runs. Baseline-subtracted. Red curve, best-fit linear filter (parametric form, gamma-Erlang function). (**B**) Saccade-locked pupil response, averaged over five easy or hard runs, for each observer. Red, hard runs. Green, easy runs. The amplitude modulation index was defined as the ratio (hard:easy) of the minima of each pair or curves. The gain modulation index was defined as the ratio (hard:easy) of the best-fit gains.

**Figure 8: Fig. S4.**
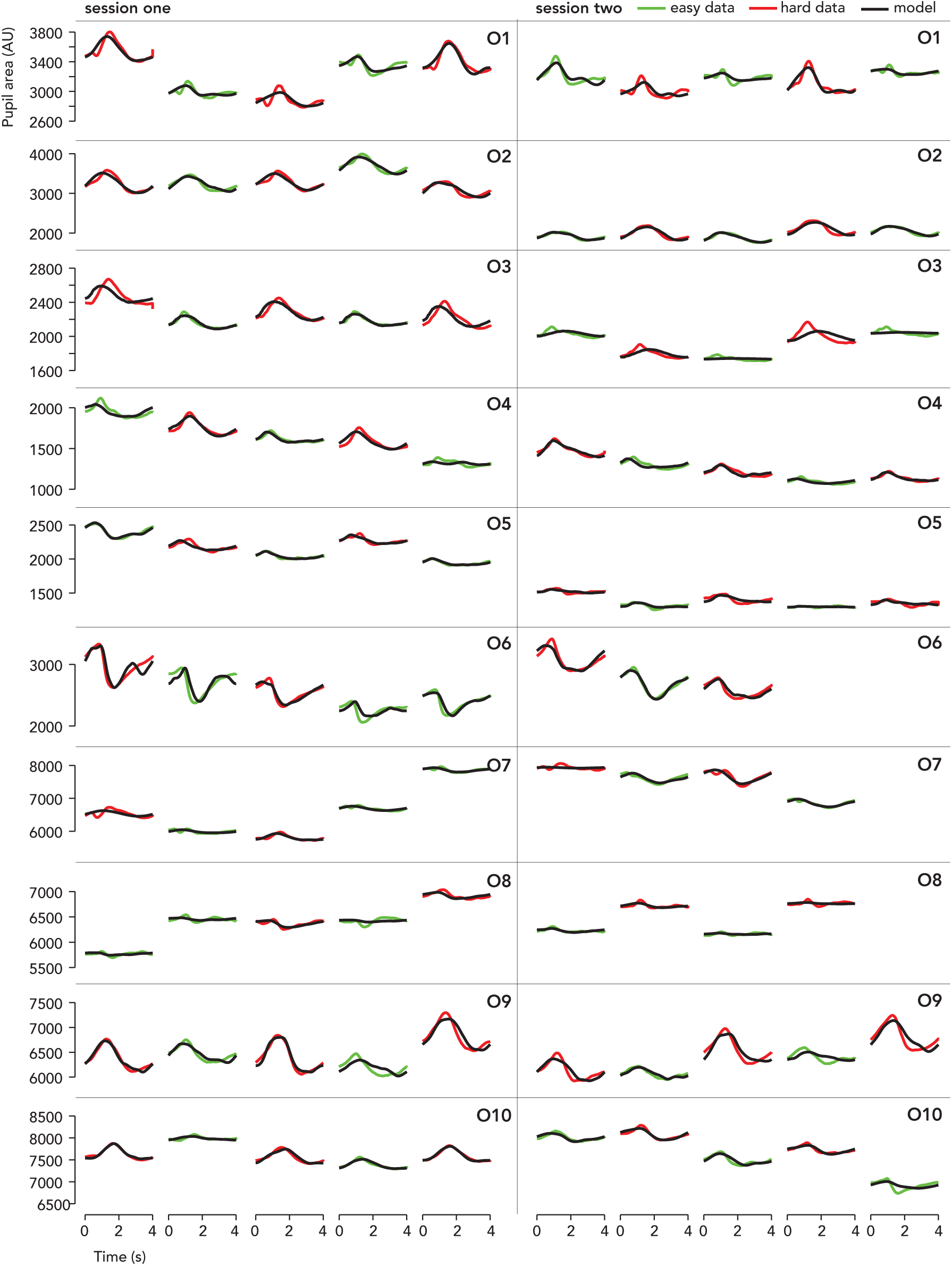
Trial-averaged pupil responses: data vs. model prediction (4 s task). Format, each column is a run and each row is an observer (O1-10). The ordinate is the same within each row to emphasize evolution of the additive offset of pupil size across runs. Colored curves, trial-averaged pupil response. Red, hard runs. Green, easy runs. Black curve, model prediction. The model captured individual differences in the shape, timing, and amplitude of the task-evoked pupil response.

**Figure 9: Fig. S5.**
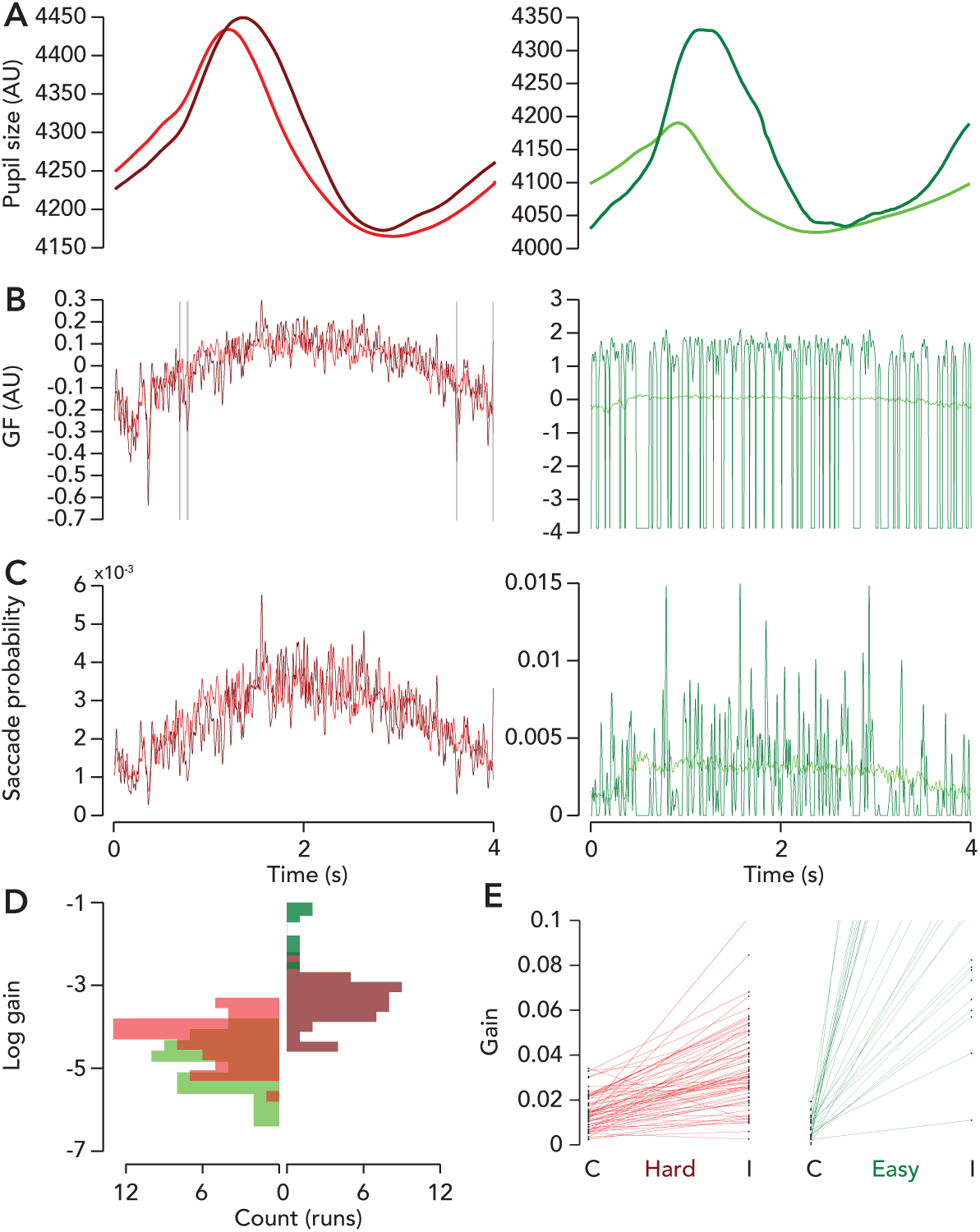
Task-evoked pupil response, generator function, saccade rate, and gain for correct and incorrect trials. (**A**) Curves, task-evoked pupil response, averaged across observers (N = 10, 4 s task). Red, hard. Green, easy. Light colors, correct trials. Dark colors, incorrect trials. Same color code used throughout. (**B**) Curves, generator function, averaged across observers. Vertical grey, times of maximal separation, i.e., d’ greater than 99% quantile of distribution of d’ over time. The difference in the generator function between hard correct and incorrect trials was minimal, with a period of maximal separation around 1 s after trial onset. The number of easy incorrect trials across all observers was very small (N trials = 48; note the y-axis limit), hence there were many time points when saccade rate was zero, leading to the large dips in the estimated easy incorrect generator function and a worse estimate overall. The sampling error was too high for easy incorrect trials to make any conclusions. (**C**) Curves, saccade rate (probability from 0 to 1), averaged across observers. High sampling error for easy incorrect (N trials = 48), as in panel B. (**D**) Best-fit (log) gain for easy and hard runs, correct and incorrect trials across all observers. Red, hard. Green, easy. Light colors (left), correct trials. Dark colors (right), incorrect trials. Only runs with *R*^2^ greater than 0.5 included (N runs = 56, easy correct; 9, easy incorrect; 62, hard correct; 46, hard incorrect; N data sets = 15; N observers = 10). Natural logarithm of gain plotted for visualization purposes due to much larger values for easy incorrect trials. (**E**) Difference in best-fit gain between correct and incorrect trials. Dots, estimated gain. C, correct. I, incorrect. Lines, difference in gain between correct and incorrect trials. Every run is shown (N runs = 145, N data sets = 15; N observers = 10); i.e., not removing trials with *R*^2^ less than 0.5, but not plotting gain values greater than 1 for visibility. It’s apparent that gain is modulated by both difficulty and accuracy.

**Figure 10: Fig. S6.**
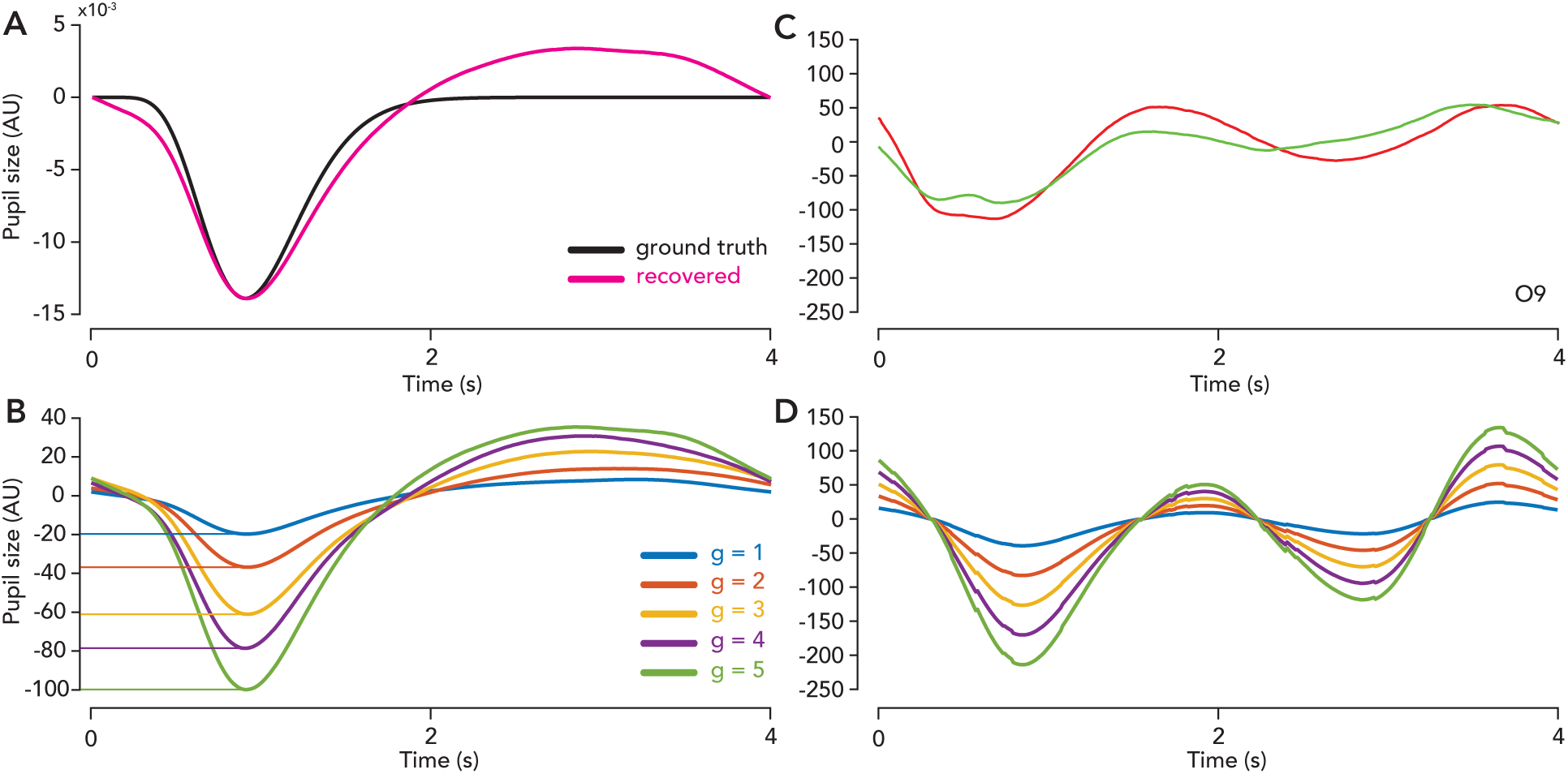
Task-dependent distortions of the saccade-locked pupil response were replicated in our model’s behavior and caused by the threshold nonlinearity. We performed a simulation to check whether our model could reproduce the observed shape of the deconvolved saccade-locked pupil response, assuming that the form of the ground truth linear filter matches our model assumption. (**A**) Black curve, ground truth linear filter used to simulate pupil data — a gamma-Erlang function. Pink curve, saccade-locked pupil response for the 4 s task, deconvolved from simulated pupil and saccade data (observer O1). (**B**) Simulated saccade-locked pupil response for the 4 s task. Different colored curves correspond to gain values ranging from 1 to 5. Horizontal lines indicate response minima, showing that they are approximately integer multiples of one another. (**C**) Run-averaged saccade-locked pupil response (data) for the 2 s task; easy and hard runs from observer O9. Green, easy. Red, hard. (**D**) Simulated saccade-locked pupil response for different gains for the 2 s task (same format as b). We used the grand-mean saccade rate function from the 2 s task from all 5 observers in the simulation. Notice the idiosyncratic distortion (multiphasic response with higher amplitude in the first trough than the second) in both the simulation (panel D) and data (panel C). That is, the model predicts the trial-duration-dependent distortion of the saccade-locked pupil response (also see Fig. S3A, O9). This simulation validates our assumption that there is a threshold non-linearity in the system. That is, if you ignore the threshold and assume that the mapping between saccades and pupil size is linear (i.e., by deconvolving the pupil response locked to saccades), you will recover a biased (distorted) estimate of the true linear filter.

**Figure 11: Fig. S7.**
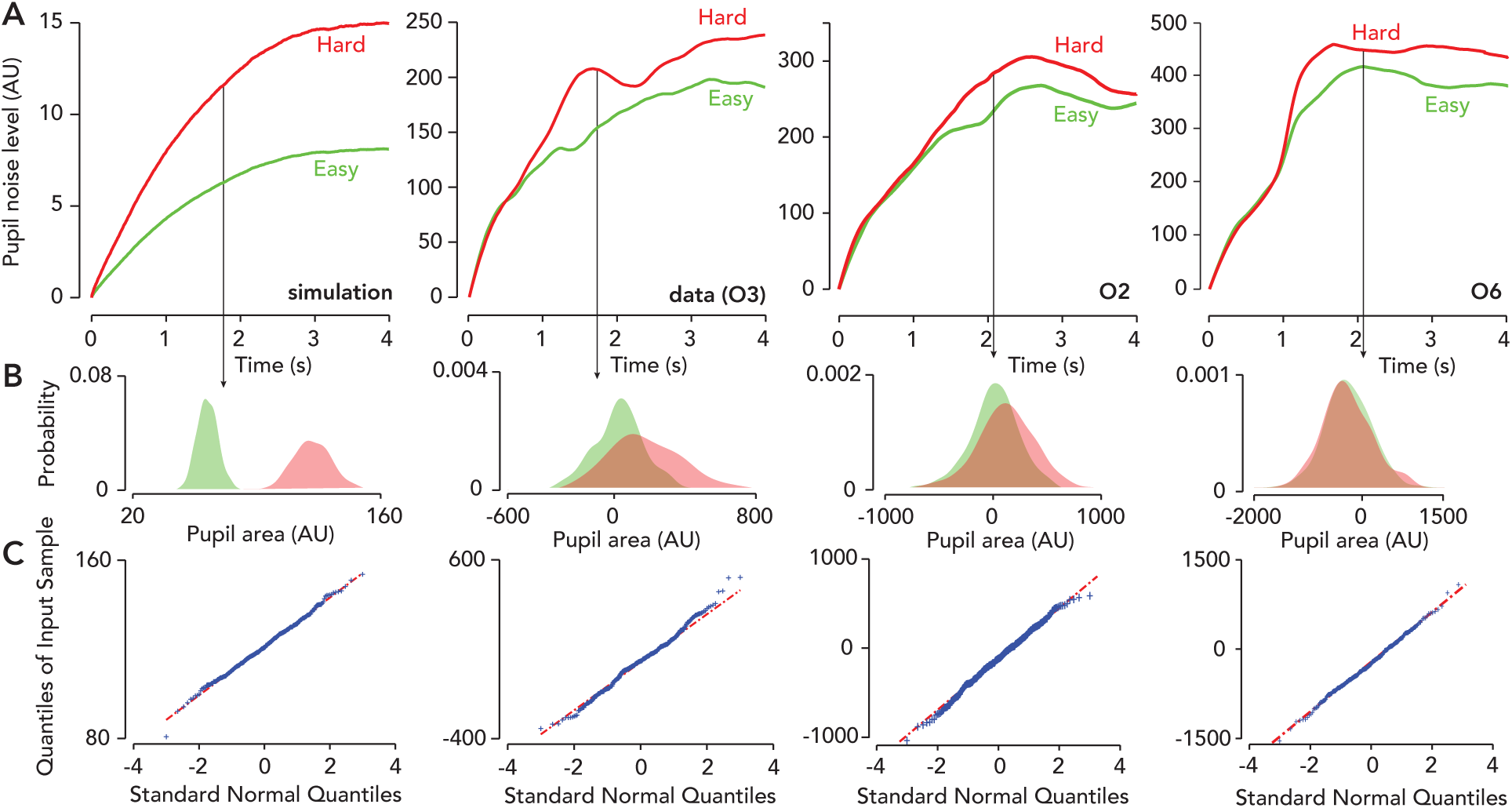
Pupil noise depends on task difficulty. (**A**) Curves, pupil noise level, quantified as the standard deviation (across trials) of the epoched task-evoked pupil response. Column 1 is a simulation and columns 2–4 are data. Red, hard runs. Green, easy runs. In our model, gain was equal to the standard deviation of pupil noise (Eq. 5) and the simulation results (column 1) matched this assumption (i.e., gain modulation [the ratio of gain, hard:easy] was equal to the best-fit scale factor that matched the easy and hard pupil noise curves). In the data (columns 2-4), the pupil noise level scaled approximately multiplicatively between hard and easy runs, and was higher for hard runs. For all three representative observers, the best-fit gain (not shown) was also larger for hard than easy runs. Epoching consisted of aligning the first time-point of each pupil response to zero. Hence, the pupil noise level was enforced to start from zero at trial onset and the non-linear increasing shape is caused by this procedure. Epoching was performed to visually emphasize scaling of the noise level. (**B**) Distribution of pupil noise across trials (N = 375) for a single time point, either 1.8 or 2.1 s. Kernel density estimate was performed using default parameters of the fitdist function in Matlab. Vertical arrow, pupil noise was measured at this time point. Compare with ref Stanten & Stark, 1966, their Fig. 4, which shows that the distribution of pupil noise is similarly modulated by both luminance and accommodation. (**C**) Q-Q plot of pupil noise distribution, for hard data only (easy looks similar), and for the same time point. Axes are standard normal quantiles vs. quantiles in input space. Data were nearly Gaussian.

**Figure 12: Fig. S8.**
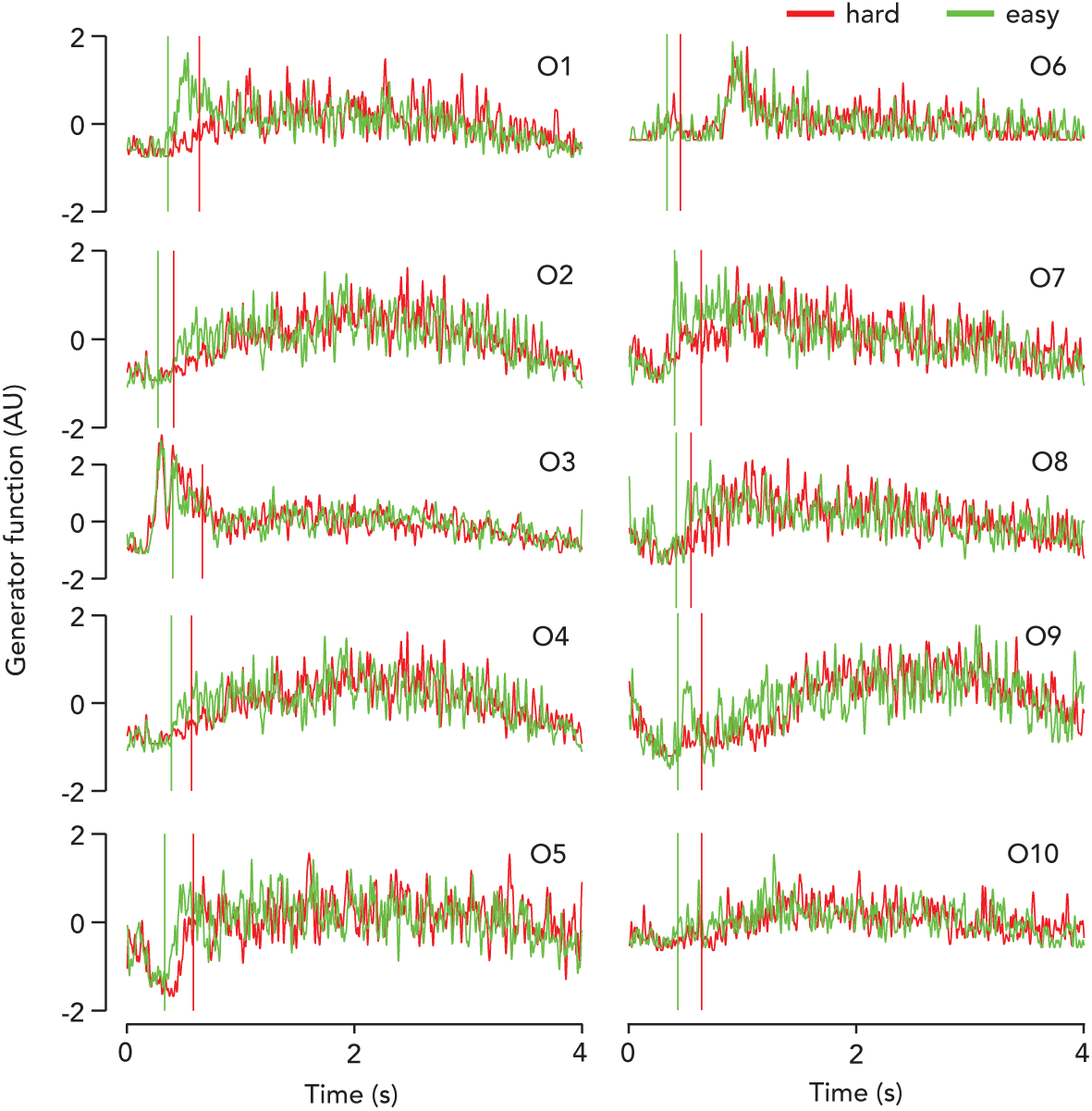
Generator functions for each observer, hard and easy runs. Curves, average generator function (trial-onset-locked) across runs (N runs = 10, N trials = 750), mean-subtracted and Gaussian-blurred over time for visualization (blurring kernel had a *σ* of 3 time samples here and throughout, wherever “Gaussian-blurred” is mentioned). Vertical lines, average reaction time. Red, hard runs. Green, easy runs.

**Figure 13: Fig. S9.**
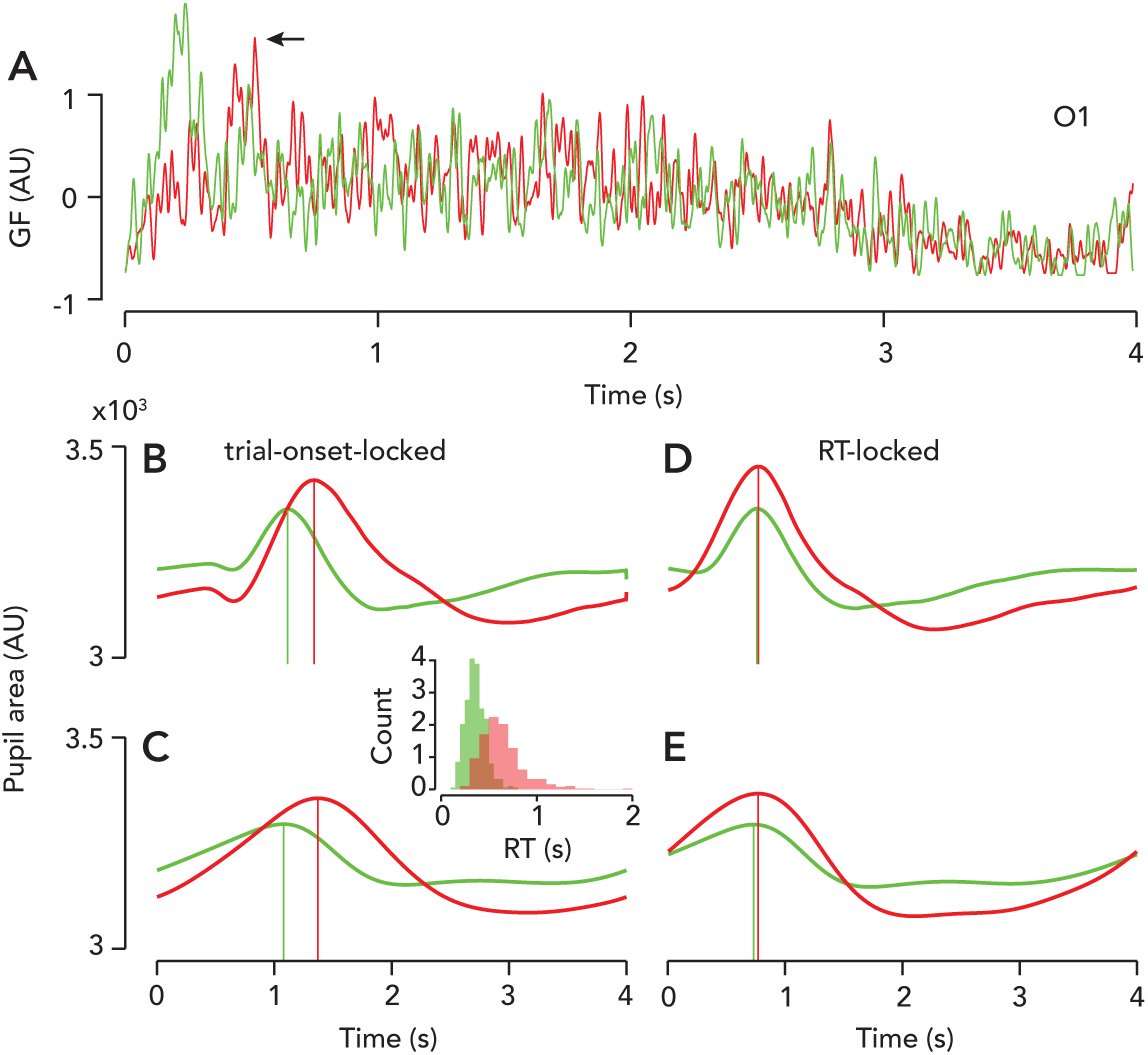
Timing of task-evoked pupil response depends on reaction-time-dependent modulations in the generator function. (**A**) Generator function time-locked to button press (“RT-locked”; N runs = 10, N trials = 750 trials). Green, easy runs. Red, hard runs. Arrow, indicating spike on hard runs that is not present in the trial-onset-locked generator function (see Fig. S8, O1). Data from observer O1. (**B**) Trial-onset locked pupil response, data. Vertical lines, time-to-peak. (**C**) Model prediction, using trial-onset-locked generator function (from Fig. S8, O1). (**D**) RT-locked pupil response. (**E**) Model prediction, using RT-locked generator function (from panel A). Inset, distribution of reaction times in seconds. Timing differences between easy and hard are abolished in model after time-locking generator function to reaction time. This demonstrates that the timing of the pupil response is driven indirectly by reaction time via modulations in the generator function (i.e., early spike for easy but not hard trials, see Fig. S8, O1).

**Figure 14: Fig. S10.**
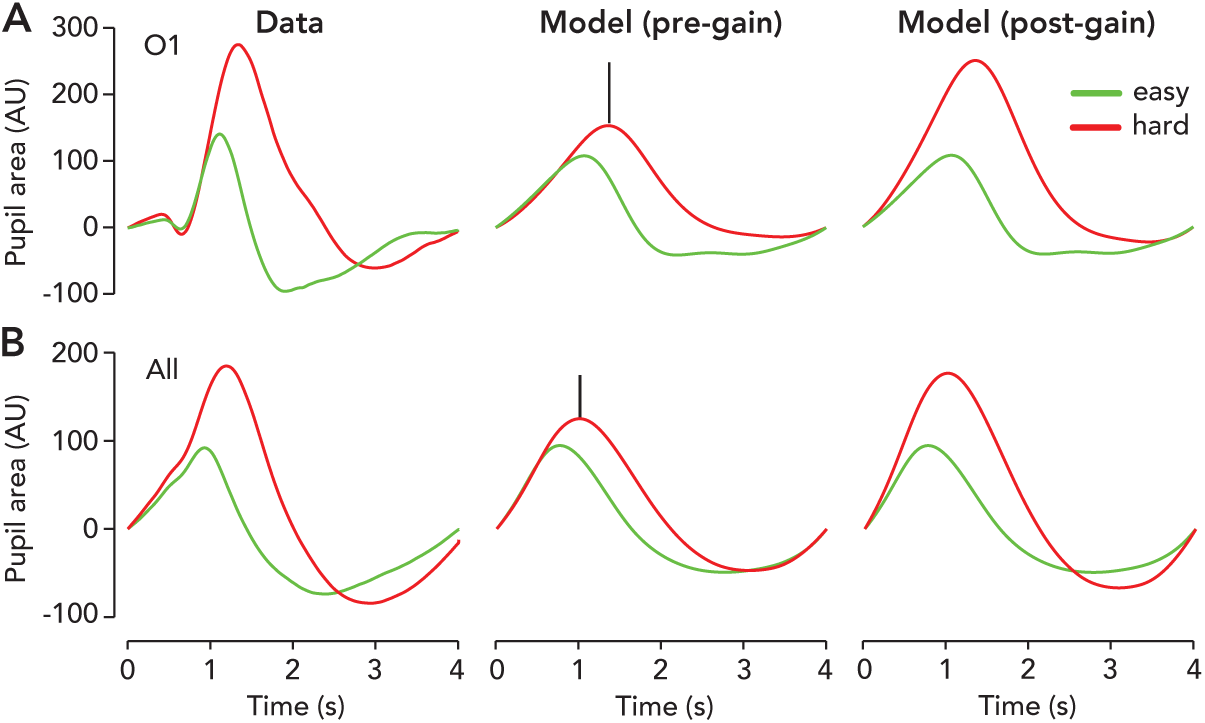
Amplitude of the task-evoked pupil response depends on generator function and gain. (**A**) Task-evoked pupil response, data and model (before and after gain modulation) for observer O1. Baseline-subtracted (i.e., pupil area at time zero is subtracted off). Red, hard trials. Green, easy trials. Left column, task-evoked pupil response, data. Center column, model prediction (pre-gain). Right column, model prediction (post-gain). Generator function was estimated separately for easy and hard trials and we only used correct trials for this analysis because accuracy also modulates pupil response amplitude. For the “pre-gain” model prediction, both curves were scaled by a single beta value from a regression between easy correct model and data, i.e., pre-gain difference in scale between correct and incorrect model predictions was preserved to emphasize amplitude modulation due to differences in generator function dynamics alone. Black vertical lines, difference between maximum of model prediction pre- vs. post-gain. This illustrates the remaining discrepancy in pupil response amplitude, i.e., the portion that is accounted for by gain. Post-gain model predictions are what we show in all other figures, i.e., gain is fit separately for easy and hard runs and used to scale each prediction. (**B**) Same format as panel A, but for the pooled data from all observers. For this analysis, we used the grand mean generator functions plotted in Fig. S5B and grand mean linear filter (N observers = 10, 4 s task). The amplitude of the task-evoked influence response is determined by both the generator function and gain, but the influence of gain is larger.

**Figure 15: Fig. S11.**
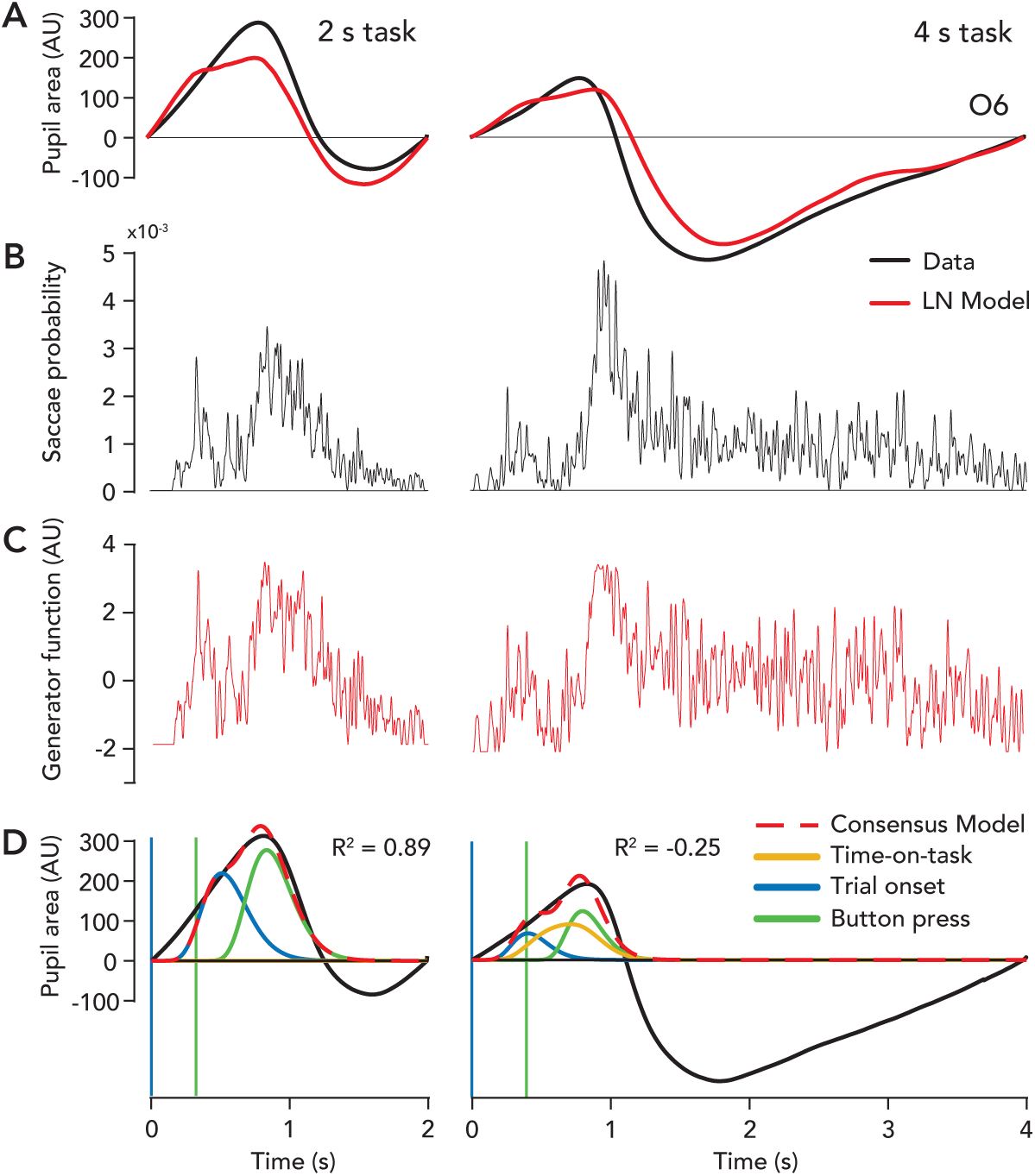
Pupil size and saccade rate were modulated by trial duration. (**A**) Curves, task-evoked pupil response (baseline-subtracted). Black, data. Red, model prediction (our model). Left, 2 s task. Right, 4 s task. Note the large pupil constriction below baseline (zero) in the 4 s, but not 2 s task. All data from a representative observer, O6. (**B**) Saccade probability for 2 and 4 s tasks (N trials = 750). (**C**) Estimated generator functions for 2 and 4 s tasks. (**D**) Consensus model fit to O6’s pupil data. Black curve, data re-plotted from panel A. Dashed red curve, model prediction (consensus model). Blue vertical line, trial onset. Green vertical line, average reaction time. Blue curve, predicted pupil response to trial-onset impulse. Green curve, predicted pupil response to button-press impulse. Yellow curve, predicted pupil response to time-on-task boxcar (i.e., whose boundaries are trial onset and reaction time). Boxcar not shown. Goodness of fit *R*^2^ reported for 2 s and 4 s tasks.

**Figure 16: Fig. S12.**
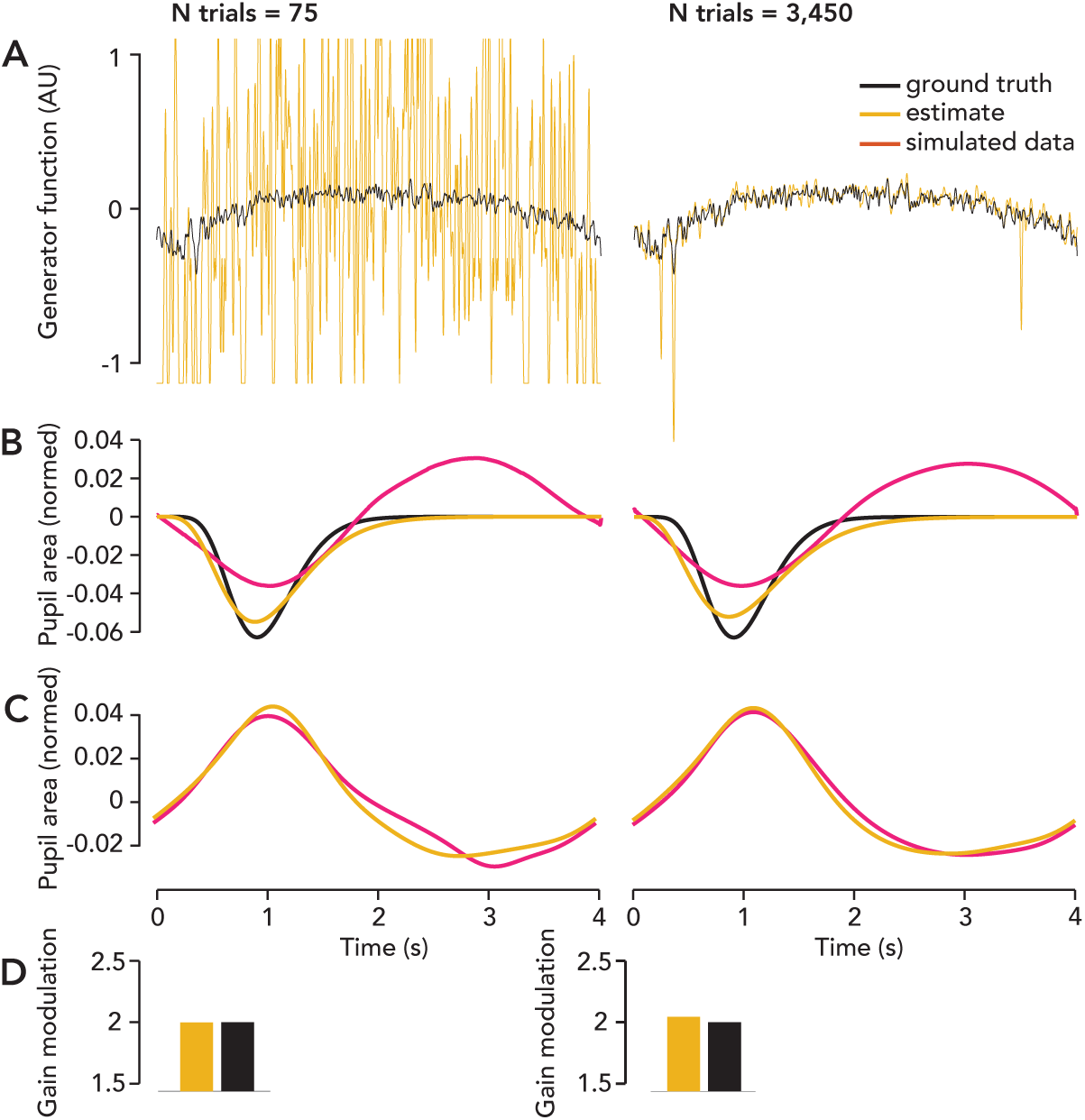
Parameter recovery. We assessed model identifiability via parameter recovery simulations wherein we asssumed a ground truth gain and generator function, simulated pupil and saccade data, and then estimated the model parameters from the synthetic data. We also varied the number of simulated trials to assess how much data is necessary to accurately recover the parameters. (**A**) Black curve, ground truth generator function (created using saccade rate from 3450 hard correct trials, N observers= 10). Yellow, estimated generator function from either 75 or 3450 simulated trials. Left column, 75 simulated trials. Right column, 3450 simulated trials. Truncated above 1.1 for yellow curve for visualization, but it continues above. All curves mean-subtracted. Estimate of generator function was accurate for 3450 trials, but not 75 trials (i.e., one run). Not shown, estimated generator function was invariant to ground truth gain. Ground truth gain equal to 1 in this panel. (**B**) Black curve, ground truth linear filter; gamma-Erlang function (tMin = 900, n = 10, f = -1). Pink curve, saccade-locked pupil response deconvolved from simulated saccade and pupil data. Yellow, estimated linear filter (parametric fit). All curves normalized to be unit vector length, i.e., the entire curve was treated as a vector and divided by its sum across elements. (**C**) Pink curve, simulated trial-averaged pupil response. Yellow curve, model prediction. *R*^2^ of fit was 96% for 75 trials and 98% for 3,450 trials. (**D**) We repeated the simulation, but increased the ground truth gain to 2. Yellow bar, ratio of estimated gain for a ground truth gain of 2 vs. 1. Gain modulation was invariant to number of trials above 75. A single run of 75 simulated trials sufficed for recovery of gain, whereas more trials were required to recover the generator function.

**Figure 17: Fig. S13.**
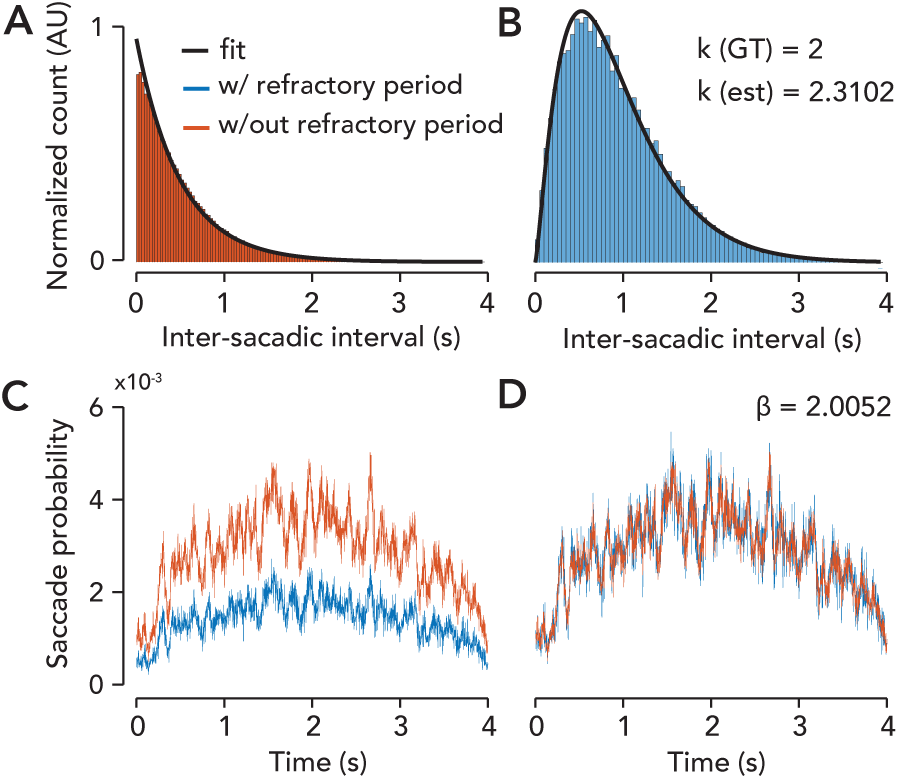
Simulation validating modelling the post-saccade refractory period with a gamma process. We generated 50000 trials of saccades from an inhomogeneous Poisson process using the average saccade rate function from observers O1–5. (**A**) Red histogram, distribution of inter-saccadic intervals (s) from the simulated Poisson process. Black curve, best-fit exponential distribution (*R*^2^ > 0.95). (**B**) Blue histogram, interval distribution after every *k*-th saccade has been been preserved and the rest have been discarded, i.e, with refractory period. Black curve, best-fit gamma distribution (*R*^2^ > 0.95). Best-fit *k* from the gamma fit is 2.31. (**C**) Saccade rate function, with and without refractory period. Same color scheme as above. (**D**) Saccade rate after scaling the two curves to match as well as possible (*R*^2^ > 0.95), i.e. outcome of a linear regression. The beta-value from the regression was 2.01 and the ground truth *k* was 2. This suggests that linear scaling correctly describes the effect of the refractory period on saccade rate, validating our procedure for inverting the refractory period in our model fitting.

**Figure 18: Fig. S14.**
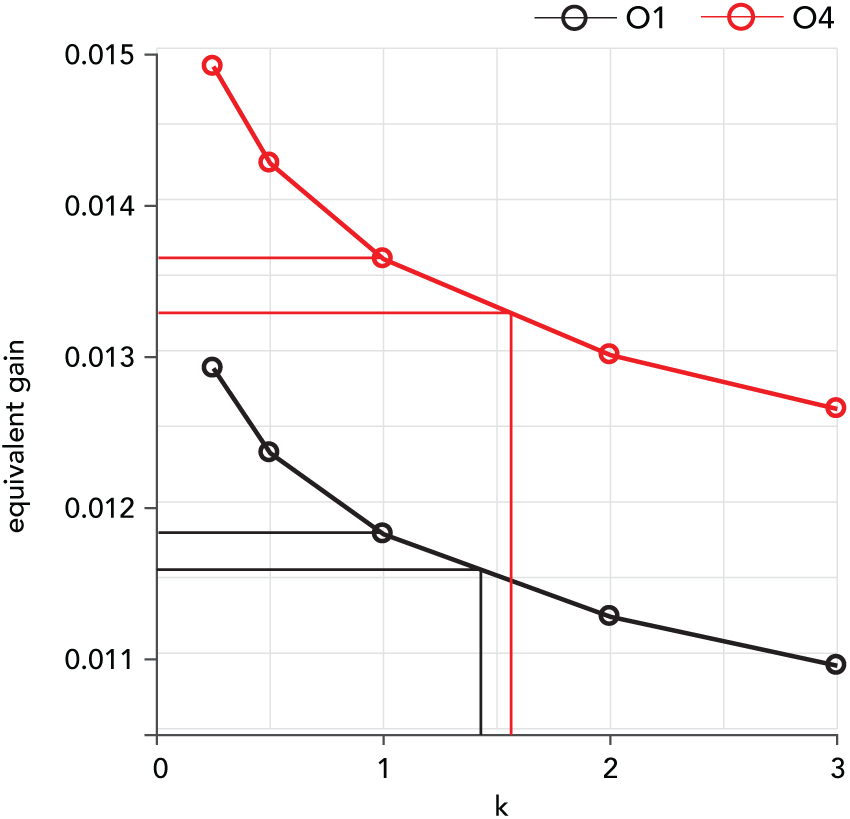
Gain traded off negligibly with *k* in the model fitting. For two example observers, we artificially varied *k*, the parameter controlling the post-saccade refractory period, and measured the change in the model’s *R*^2^ and best-fit gain. Abscissa, *k* was set to 0.25, 0.5, 1, 2, or 3. Each dot represents a *k* of one of these values. Vertical lines represent best-fit *k* value for that observer. Ordinate, solution for gain corresponding to the chosen *k*. Horizontal lines, best-fit gain corresponding to the best-fit *k* or a *k* of 1; to emphasize the very small change in gain when the refractory period is removed from the model (i.e., when *k* is set to 1). Black, for observer O1. Red, for observer O4. Compare the distance between the horizontal lines with the difference in the best-fit gain for hard vs. easy runs of trials, which was more than a order of magnitude larger. This demonstrates that *k* negligibly influences the solution for gain. Not shown, difference in *R*^2^ between the different points on the graph was always less than 0.0001, demonstrating the refractory period’s inclusion in the model did not influence the model’s goodness of fit.

**Table 1:** Best-fit model parameters, goodness of fit, and data summary metrics for every run and every observer. Each row corresponds with a run of 75 trials. “Data sets 11-15” correspond to the data from 2 s task for Observers O6-10, so data set “11” in the table is actually observer O6, 2 s task. Data sets 1-10 correspond to the 4 s task for observers O1-10. For “hard or easy,” 1 is hard and 0 is easy. “Reaction time” is in seconds.

| Data set | gain | offset | $k$ | hard or easy | session | accuracy | reaction time | $R^2$ |
| --- | --- | --- | --- | --- | --- | --- | --- | --- |
| 1 | 0.0190 | 3543.2711 | 1.4424 | 1 | 2 | 0.7600 | 0.6032 | 0.7944 |
| 1 | 0.0079 | 2989.0147 | 1.4424 | 0 | 2 | 0.9600 | 0.3613 | 0.5694 |
| 1 | 0.0096 | 2882.0478 | 1.4424 | 1 | 2 | 0.6933 | 0.6326 | 0.6502 |
| 1 | 0.0140 | 3345.3563 | 1.4424 | 0 | 2 | 0.9867 | 0.3616 | 0.7570 |
| 1 | 0.0211 | 3407.3333 | 1.4424 | 1 | 2 | 0.6800 | 0.7431 | 0.8556 |
| 1 | 0.0218 | 3209.8797 | 1.4424 | 0 | 1 | 0.9733 | 0.3367 | 0.7573 |
| 1 | 0.0155 | 3003.4486 | 1.4424 | 1 | 1 | 0.8000 | 0.6221 | 0.5438 |
| 1 | 0.0063 | 3183.8036 | 1.4424 | 0 | 1 | 1.0000 | 0.3776 | 0.4984 |
| 1 | 0.0208 | 3100.1904 | 1.4424 | 1 | 1 | 0.7600 | 0.5845 | 0.8449 |
| 1 | 0.0050 | 3258.1470 | 1.4424 | 0 | 1 | 0.9733 | 0.3631 | 0.6719 |
| 2 | 0.0109 | 3248.2887 | 0.8659 | 1 | 1 | 0.7600 | 0.4382 | 0.8032 |
| 2 | 0.0156 | 3229.7041 | 0.8659 | 0 | 1 | 0.8933 | 0.2672 | 0.9207 |
| 2 | 0.0080 | 3276.1298 | 0.8659 | 1 | 1 | 0.7200 | 0.4483 | 0.8021 |
| 2 | 0.0079 | 3699.5384 | 0.8659 | 1 | 1 | 0.7333 | 0.4109 | 0.8753 |
| 2 | 0.0089 | 3088.8217 | 0.8659 | 0 | 1 | 1.0000 | 0.2703 | 0.8823 |
| 2 | 0.0055 | 1918.6730 | 0.8659 | 0 | 2 | 0.9867 | 0.2586 | 0.9236 |
| 2 | 0.0076 | 1986.9199 | 0.8659 | 1 | 2 | 0.6933 | 0.3635 | 0.9392 |
| 2 | 0.0056 | 1879.7111 | 0.8659 | 0 | 2 | 1.0000 | 0.2958 | 0.9127 |
| 2 | 0.0079 | 2108.8628 | 0.8659 | 1 | 2 | 0.7467 | 0.3944 | 0.8760 |
| 2 | 0.0054 | 2051.9358 | 0.8659 | 0 | 2 | 1.0000 | 0.2733 | 0.9403 |
| 3 | 0.0184 | 2474.6711 | 1.2568 | 1 | 2 | 0.7333 | 0.7498 | 0.4619 |
| 3 | 0.0148 | 2152.3261 | 1.2568 | 0 | 2 | 1.0000 | 0.4787 | 0.9056 |
| 3 | 0.0156 | 2287.1225 | 1.2568 | 1 | 2 | 0.7200 | 0.7275 | 0.8366 |
| 3 | 0.0161 | 2178.3422 | 1.2568 | 0 | 2 | 0.9867 | 0.4556 | 0.9360 |
| 3 | 0.0198 | 2212.8509 | 1.2568 | 1 | 2 | 0.7200 | 0.6607 | 0.6828 |
| 3 | 0.0061 | 2032.0018 | 1.2568 | 0 | 1 | 0.9733 | 0.3768 | 0.3413 |
| 3 | 0.0101 | 1798.8518 | 1.2568 | 1 | 1 | 0.8000 | 0.5775 | 0.4600 |
| 3 | 0.0008 | 1738.5184 | 1.2568 | 0 | 1 | 0.9733 | 0.3511 | 0.0140 |
| 3 | 0.0079 | 2007.2994 | 1.2568 | 1 | 1 | 0.7867 | 0.5968 | 0.2332 |
| 3 | 0.0009 | 2042.7166 | 1.2568 | 0 | 1 | 1.0000 | 0.3515 | 0.0130 |
| 4 | 0.0155 | 1951.8998 | 1.5471 | 0 | 1 | 0.9867 | 0.3557 | 0.5576 |
| 4 | 0.0160 | 1756.9504 | 1.5471 | 1 | 1 | 0.7200 | 0.6700 | 0.9019 |
| 4 | 0.0157 | 1615.9255 | 1.5471 | 0 | 1 | 0.9733 | 0.4040 | 0.8490 |
| 4 | 0.0119 | 1581.1618 | 1.5471 | 1 | 1 | 0.7067 | 0.6268 | 0.7659 |
| 4 | 0.0063 | 1318.6639 | 1.5471 | 0 | 1 | 0.9600 | 0.5331 | 0.1063 |
| 4 | 0.0242 | 1480.6169 | 1.5471 | 1 | 2 | 0.8000 | 0.4916 | 0.9178 |
| 4 | 0.0121 | 1300.0452 | 1.5471 | 0 | 2 | 0.9867 | 0.3257 | 0.5589 |
| 4 | 0.0160 | 1213.0523 | 1.5471 | 1 | 2 | 0.7467 | 0.5244 | 0.7273 |
| 4 | 0.0044 | 1093.4174 | 1.5471 | 0 | 2 | 1.0000 | 0.3351 | 0.5030 |
| 4 | 0.0099 | 1139.6118 | 1.5471 | 1 | 2 | 0.7067 | 0.5222 | 0.8921 |
| 5 | 0.0237 | 2398.5941 | 2.4954 | 0 | 2 | 0.9867 | 0.3068 | 0.9164 |
| 5 | 0.0128 | 2179.9517 | 2.4954 | 1 | 2 | 0.7867 | 0.5506 | 0.5186 |
| 5 | 0.0104 | 2037.3889 | 2.4954 | 0 | 2 | 1.0000 | 0.2688 | 0.9309 |
| 5 | 0.0130 | 2271.0164 | 2.4954 | 1 | 2 | 0.6933 | 0.5875 | 0.7102 |
| 5 | 0.0101 | 1941.0339 | 2.4954 | 0 | 2 | 0.9867 | 0.2756 | 0.9390 |
| 5 | 0.0075 | 1520.6753 | 2.4954 | 1 | 1 | 0.7467 | 0.5937 | 0.4654 |
| 5 | 0.0106 | 1312.8609 | 2.4954 | 0 | 1 | 1.0000 | 0.3837 | 0.6769 |
| 5 | 0.0117 | 1404.7837 | 2.4954 | 1 | 1 | 0.7067 | 0.5995 | 0.5811 |
| 5 | 0.0039 | 1297.5804 | 2.4954 | 0 | 1 | 1.0000 | 0.4254 | 0.2323 |
| 5 | 0.0082 | 1351.8020 | 2.4954 | 1 | 1 | 0.7333 | 0.5842 | 0.2989 |
| 6 | 0.0175 | 2965.4310 | 1.0131 | 1 | 2 | 0.7600 | 0.4805 | 0.8345 |
| 6 | 0.0184 | 2697.7299 | 1.0131 | 0 | 2 | 0.9867 | 0.3511 | 0.6839 |
| 6 | 0.0081 | 2531.9243 | 1.0131 | 1 | 2 | 0.6667 | 0.4779 | 0.9062 |
| 6 | 0.0100 | 2252.9978 | 1.0131 | 0 | 2 | 0.9867 | 0.3596 | 0.4257 |
| 6 | 0.0083 | 2386.2589 | 1.0131 | 0 | 1 | 1.0000 | 0.3282 | 0.8539 |
| 6 | 0.0106 | 3073.5765 | 1.0131 | 1 | 1 | 0.8133 | 0.4383 | 0.8515 |
| 6 | 0.0120 | 2679.3602 | 1.0131 | 0 | 1 | 0.9733 | 0.2900 | 0.9689 |
| 6 | 0.0102 | 2572.7380 | 1.0131 | 1 | 1 | 0.6933 | 0.4045 | 0.8170 |
| 7 | 0.0021 | 6538.0594 | 1.0604 | 1 | 1 | 0.7467 | 0.6269 | 0.2482 |
| 7 | 0.0015 | 5988.1508 | 1.0604 | 0 | 1 | 0.9867 | 0.4093 | 0.5716 |
| 7 | 0.0041 | 5805.8896 | 1.0604 | 1 | 1 | 0.7600 | 0.7228 | 0.8303 |
| 7 | 0.0019 | 6687.6094 | 1.0604 | 0 | 1 | 1.0000 | 0.4082 | 0.6395 |
| 7 | 0.0029 | 7860.2022 | 1.0604 | 0 | 2 | 1.0000 | 0.3838 | 0.7781 |
| 7 | 0.0001 | 7930.7733 | 1.0604 | 1 | 2 | 0.6800 | 0.6150 | 0.0006 |
| 7 | 0.0058 | 7602.7974 | 1.0604 | 0 | 2 | 1.0000 | 0.4331 | 0.8001 |
| 7 | 0.0067 | 7649.7935 | 1.0604 | 1 | 2 | 0.7067 | 0.5726 | 0.8466 |
| 7 | 0.0046 | 6848.6961 | 1.0604 | 0 | 2 | 1.0000 | 0.3655 | 0.8940 |
| 8 | 0.0036 | 5769.1064 | 0.9565 | 0 | 1 | 0.9867 | 0.4076 | 0.2848 |
| 8 | 0.0029 | 6454.8527 | 0.9565 | 0 | 1 | 0.9733 | 0.4055 | 0.1843 |
| 8 | 0.0054 | 6362.5391 | 0.9565 | 1 | 1 | 0.7067 | 0.4966 | 0.7217 |
| 8 | 0.0023 | 6420.5245 | 0.9565 | 0 | 1 | 0.9867 | 0.4228 | 0.0573 |
| 8 | 0.0046 | 6918.0012 | 0.9565 | 1 | 2 | 0.7867 | 0.4875 | 0.5238 |
| 8 | 0.0029 | 6228.6349 | 0.9565 | 0 | 2 | 0.9600 | 0.3946 | 0.6410 |
| 8 | 0.0027 | 6719.2112 | 0.9565 | 1 | 2 | 0.6667 | 0.5846 | 0.5572 |
| 8 | 0.0019 | 6162.6933 | 0.9565 | 0 | 2 | 0.9867 | 0.4444 | 0.1543 |
| 8 | 0.0014 | 6766.5297 | 0.9565 | 1 | 2 | 0.7333 | 0.6306 | 0.0778 |
| 9 | 0.0223 | 6375.5461 | 0.9442 | 1 | 1 | 0.7733 | 0.5499 | 0.9324 |
| 9 | 0.0144 | 6463.7846 | 0.9442 | 0 | 1 | 0.9733 | 0.4187 | 0.8261 |
| 9 | 0.0233 | 6393.1706 | 0.9442 | 1 | 1 | 0.7467 | 0.6281 | 0.9607 |
| 9 | 0.0132 | 6189.4111 | 0.9442 | 0 | 1 | 0.9733 | 0.4532 | 0.5639 |
| 9 | 0.0212 | 6839.2854 | 0.9442 | 1 | 1 | 0.6667 | 0.6247 | 0.8879 |
| 9 | 0.0145 | 6131.4102 | 0.9442 | 1 | 2 | 0.7333 | 0.5684 | 0.8147 |
| 9 | 0.0098 | 6069.4518 | 0.9442 | 0 | 2 | 0.9867 | 0.4162 | 0.8717 |
| 9 | 0.0192 | 6551.2339 | 0.9442 | 1 | 2 | 0.7067 | 0.6361 | 0.8634 |
| 9 | 0.0098 | 6407.0855 | 0.9442 | 0 | 2 | 1.0000 | 0.4212 | 0.4057 |
| 9 | 0.0186 | 6808.4108 | 0.9442 | 1 | 2 | 0.7600 | 0.5346 | 0.8372 |
| 10 | 0.0209 | 7639.3860 | 0.8506 | 1 | 1 | 0.7333 | 0.6881 | 0.9491 |
| 10 | 0.0049 | 7989.0090 | 0.8506 | 0 | 1 | 1.0000 | 0.4196 | 0.4359 |
| 10 | 0.0192 | 7560.6919 | 0.8506 | 1 | 1 | 0.7067 | 0.6191 | 0.9100 |
| 10 | 0.0110 | 7388.8304 | 0.8506 | 0 | 1 | 1.0000 | 0.4366 | 0.9462 |
| 10 | 0.0259 | 7600.1529 | 0.8506 | 1 | 1 | 0.7333 | 0.6572 | 0.9397 |
| 10 | 0.0052 | 8009.4760 | 0.8506 | 0 | 2 | 0.9867 | 0.4198 | 0.6688 |
| 10 | 0.0058 | 8080.9123 | 0.8506 | 1 | 2 | 0.6667 | 0.6161 | 0.8562 |
| 10 | 0.0122 | 7492.2008 | 0.8506 | 0 | 2 | 0.9867 | 0.4260 | 0.8467 |
| 10 | 0.0051 | 7727.8971 | 0.8506 | 1 | 2 | 0.7200 | 0.6047 | 0.8375 |
| 10 | 0.0063 | 6916.7567 | 0.8506 | 0 | 2 | 0.9867 | 0.4436 | 0.4214 |
| 11 | 0.0100 | 2979.6322 | 1.1329 | 1 | 2 | 0.7333 | 0.2981 | 0.4117 |
| 11 | 0.0121 | 2790.8970 | 1.1329 | 0 | 2 | 0.8800 | 0.2959 | 0.9588 |
| 11 | 0.0197 | 2877.8471 | 1.1329 | 1 | 2 | 0.7733 | 0.3266 | 0.9295 |
| 11 | 0.0104 | 2962.4148 | 1.1329 | 0 | 2 | 0.9067 | 0.2956 | 0.7834 |
| 11 | 0.0084 | 2451.5970 | 1.1329 | 0 | 1 | 0.9200 | 0.3888 | 0.7558 |
| 11 | 0.0148 | 2703.3049 | 1.1329 | 1 | 1 | 0.8133 | 0.3557 | 0.9148 |
| 11 | 0.0098 | 2786.4994 | 1.1329 | 0 | 1 | 0.9600 | 0.2771 | 0.9475 |
| 11 | 0.0121 | 2902.6339 | 1.1329 | 1 | 1 | 0.7733 | 0.3442 | 0.9501 |
| 12 | 0.0077 | 7269.4646 | 1.2817 | 1 | 1 | 0.7200 | 0.4754 | 0.8117 |
| 12 | 0.0049 | 7375.6144 | 1.2817 | 0 | 1 | 0.9867 | 0.3290 | 0.8309 |
| 12 | 0.0037 | 7403.0022 | 1.2817 | 0 | 1 | 0.9867 | 0.3040 | 0.6500 |
| 12 | 0.0030 | 6949.2244 | 1.2817 | 1 | 1 | 0.7200 | 0.4978 | 0.2411 |
| 12 | 0.0047 | 6965.3848 | 1.2817 | 1 | 2 | 0.7867 | 0.4621 | 0.5653 |
| 12 | 0.0030 | 6523.1524 | 1.2817 | 0 | 2 | 0.9867 | 0.2580 | 0.4742 |
| 12 | 0.0049 | 6631.0211 | 1.2817 | 1 | 2 | 0.7467 | 0.4834 | 0.4547 |
| 12 | 0.0054 | 7760.6452 | 1.2817 | 0 | 2 | 1.0000 | 0.3372 | 0.8281 |
| 13 | 0.0064 | 6663.5038 | 0.9315 | 0 | 1 | 0.9733 | 0.3395 | 0.9049 |
| 13 | 0.0046 | 6739.4979 | 0.9315 | 1 | 1 | 0.7867 | 0.4513 | 0.7211 |
| 13 | 0.0042 | 6288.4910 | 0.9315 | 0 | 1 | 0.9733 | 0.3467 | 0.8858 |
| 13 | 0.0059 | 7213.5414 | 0.9315 | 1 | 1 | 0.6133 | 0.4674 | 0.7170 |
| 13 | 0.0045 | 6638.4800 | 0.9315 | 0 | 1 | 0.9600 | 0.3084 | 0.7296 |
| 13 | 0.0028 | 6692.1295 | 0.9315 | 1 | 2 | 0.6933 | 0.5154 | 0.2187 |
| 13 | 0.0031 | 5956.9870 | 0.9315 | 0 | 2 | 0.9200 | 0.4098 | 0.5333 |
| 13 | 0.0045 | 6492.0587 | 0.9315 | 1 | 2 | 0.7333 | 0.5261 | 0.8750 |
| 13 | 0.0029 | 6633.6915 | 0.9315 | 0 | 2 | 0.9867 | 0.3036 | 0.7843 |
| 13 | 0.0060 | 6579.1206 | 0.9315 | 1 | 2 | 0.8133 | 0.5370 | 0.9377 |
| 13 | 0.0048 | 6574.9278 | 0.9315 | 0 | 2 | 0.9733 | 0.3395 | 0.8458 |
| 14 | 0.0074 | 6657.7994 | 0.9002 | 0 | 1 | 0.9600 | 0.3232 | 0.4250 |
| 14 | 0.0121 | 7107.0955 | 0.9002 | 1 | 1 | 0.7733 | 0.4666 | 0.9461 |
| 14 | 0.0124 | 7005.3981 | 0.9002 | 0 | 1 | 0.9467 | 0.2818 | 0.7621 |
| 14 | 0.0120 | 7489.9352 | 0.9002 | 1 | 1 | 0.7600 | 0.4720 | 0.9656 |
| 14 | 0.0115 | 6931.3348 | 0.9002 | 0 | 1 | 0.9600 | 0.3400 | 0.9324 |
| 14 | 0.0112 | 6599.2860 | 0.9002 | 1 | 2 | 0.7600 | 0.4809 | 0.9771 |
| 14 | 0.0071 | 6680.3687 | 0.9002 | 0 | 2 | 0.9600 | 0.2984 | 0.8889 |
| 14 | 0.0100 | 7081.5785 | 0.9002 | 1 | 2 | 0.6933 | 0.5368 | 0.9623 |
| 14 | 0.0059 | 6637.5857 | 0.9002 | 0 | 2 | 0.9867 | 0.3193 | 0.8627 |
| 14 | 0.0121 | 6602.6390 | 0.9002 | 1 | 2 | 0.7067 | 0.4300 | 0.9684 |
| 15 | 0.0213 | 7186.0319 | 0.9331 | 1 | 1 | 0.6533 | 0.6158 | 0.7207 |
| 15 | 0.0157 | 7609.7498 | 0.9331 | 0 | 1 | 0.9867 | 0.3839 | 0.9292 |
| 15 | 0.0194 | 7720.3537 | 0.9331 | 1 | 1 | 0.6533 | 0.5924 | 0.9007 |
| 15 | 0.0139 | 7259.8610 | 0.9331 | 0 | 1 | 1.0000 | 0.4114 | 0.7395 |
| 15 | 0.0162 | 6950.6565 | 0.9331 | 1 | 1 | 0.6533 | 0.5693 | 0.7176 |
| 15 | 0.0194 | 7817.6803 | 0.9331 | 0 | 2 | 1.0000 | 0.3524 | 0.9128 |
| 15 | 0.0120 | 8019.8782 | 0.9331 | 1 | 2 | 0.7067 | 0.5227 | 0.9209 |
| 15 | 0.0158 | 7411.1037 | 0.9331 | 0 | 2 | 0.9867 | 0.3541 | 0.9600 |
| 15 | 0.0157 | 8011.6845 | 0.9331 | 1 | 2 | 0.7067 | 0.5047 | 0.9216 |
| 15 | 0.0081 | 8582.1398 | 0.9331 | 0 | 2 | 1.0000 | 0.3860 | 0.9412 |

